# Genomic Characterization of Posttraumatic Stress Disorder in a Large US Military Veteran Sample

**DOI:** 10.1101/764001

**Authors:** Murray B. Stein, Daniel F. Levey, Zhongshan Cheng, Frank R. Wendt, Kelly Harrington, Kelly Cho, Rachel Quaden, Krishnan Radhakrishnan, Matthew J. Girgenti, Yuk-Lam Anne Ho, Daniel Posner, PTSD Working Group of the Psychiatric Genomics Consortium (PGC), Traumatic Stress Brain Research Study Group, VA Million Veteran Program, VA Cooperative Studies Program, Mihaela Aslan, Ronald S. Duman, Hongyu Zhao, Renato Polimanti, John Concato, Joel Gelernter

## Abstract

Individuals vary in their liability to develop Posttraumatic Stress Disorder (PTSD), the symptoms of which are highly heterogeneous, following exposure to life-threatening trauma. Understanding genetic factors that contribute to the biology of PTSD is critical for refining diagnosis and developing new treatments. Using genetic data from more than 250,000 participants in the Million Veteran Program, genomewide association analyses were conducted using a validated electronic health record-based algorithmically-defined PTSD diagnosis phenotype (48,221 cases and 217,223 controls), and PTSD quantitative symptom phenotypes (212,007 individuals). We identified several genome-wide significant loci in the case-control analyses, and numerous such loci in the quantitative trait analyses, including some (e.g., *MAD1L1*; *TCF4; CRHR1*) that were associated with multiple symptom sub-domains and total symptom score, and others that were more specific to certain symptom sub-domains (e.g., *CAMKV* to re-experiencing; *SOX6* to hyperarousal). Genetic correlations between all pairs of symptom sub-domains and their total were very high (r_g_ 0.93 – 0.98) supporting validity of the PTSD diagnostic construct. We also demonstrate strong shared heritability with a range of traits, show that heritability persists when conditioned on other major psychiatric disorders, present independent replication results, provide support for one of the implicated genes in postmortem brain of individuals with PTSD, and use this information to identify potential drug repositioning candidates. These results point to the utility of genetics to inform and validate the biological coherence of the PTSD syndrome despite considerable heterogeneity at the symptom level, and to provide new directions for treatment development.

## INTRODUCTION

Posttraumatic stress disorder (PTSD) is a common mental disorder that can occur after exposure to extreme, life threating stress (American Psychiatric Association, 2013). Although most Americans (50-85%) experience traumatic events over a lifetime, most do not develop PTSD – lifetime PTSD prevalence is approximately 7% (Kessler and Wang, 2008), suggesting differential resilience to stress and vulnerability to the disorder (Atwoli et al., 2015). There is a substantial heritable basis for PTSD risk (Stein et al., 2002; Wolf et al., 2018), and evidence from genome-wide association studies (GWAS) shows that PTSD, like other mental disorders (Sullivan et al., 2018), is highly polygenic (Benjet et al., 2016; Daskalakis et al., 2018; Duncan et al., 2018; Nievergelt et al., 2018; Stein et al., 2016; Xie et al., 2013).

PTSD symptoms vary widely among individuals, with the current DSM-5 definition permitting up to 163,120 unique conformations for assembly of the disorder (Galatzer-Levy and Bryant, 2013). Recognizing that this phenotypic heterogeneity may impair the detection of genetic risk factors (Stein, 2018), alternate phenotypes or sub-phenotypes that may reflect biologically more homogeneous entities have been examined. One such approach that we have used is to focus on a core component of the PTSD phenotype, re-experiencing (in DSM-IV, and referred to in DSM-5 as intrusion) symptoms, which has revealed numerous genome-wide significant risk loci, including some that had long been *a priori* functional candidates (e.g., *CRHR1*) (Gelernter et al., 2019). An additional advantage to looking at sub-phenotypes is the ability to measure them continuously, enabling the incorporation of more information and larger informative sample sizes (and therefore greater power) than is possible with a case-control design. Other approaches to surmounting phenotypic heterogeneity include the accrual of ever larger sample sizes, which has proven effective for other mental disorders such as schizophrenia (Sullivan et al., 2018) and major depression (Wray et al., 2018). The largest PTSD GWAS study to date, conducted by the PTSD Working Group of the Psychiatric Genomics Consortium (PGC-PTSD) included approximately 30,000 cases (Nievergelt et al., 2018), still moderately-powered by contemporary GWAS standards, assimilating information from over 60 separate heterogeneous sources. The use of biobanks with relatively large numbers of PTSD cases offers the opportunity to provide unprecedented sample size, ascertain granularity of symptoms and associated comorbid conditions and, importantly, uniformity of phenotypic and genotypic platforms (Radhakrishnan et al., 2019).

The current investigation was conducted within the US Veterans Affairs Million Veteran Program (MVP) (Gaziano et al., 2016) and included several PTSD phenotypic definitions: a validated, algorithmically-defined [binary] case-control approach using data from the electronic health record (EHR); and a quantitative trait approach encompassing sub-phenotypes based on recent self-reported symptoms: re-experiencing (in an expanded sample from that previously reported (Gelernter et al., 2019)), avoidance, hyperarousal and a total index of recent symptom severity (PCL total). Heritability of each of these phenotypes as well as observed (phenotypic) and genetic (r_g_) correlations were examined with the aim of determining coherence among them; r_g_ with other behavioral and health-related traits were also examined. Results for the phenotype with greatest estimated heritability in MVP were replicated in an external dataset using polygenic risk scores (PRSs) (Wray et al., 2014) from MVP-PTSD as predictors of PTSD case-control status in PGC-PTSD Freeze 2.0 (Nievergelt et al., 2018). Postmortem brain expression analysis (Girgenti and Duman, 2018) was subsequently used to validate key GWAS observations. Additional post-GWAS analyses included enrichment (tissue; pathway) and transcriptome-wide (Gamazon et al., 2015) analyses. Phenome-wide analysis (PheWAS) was conducted for the top GWAS hits against the VA-MVP EHR (Bush et al., 2016; Smoller, 2018) and possible drug candidates for repositioning were identified using publicly available drug-genomic databases.

Aims of these analyses are to provide (1) the largest single-source GWAS of PTSD to date with the greatest multi-ethnic representation, (2) with replication of key associations in other datasets, along with ascertaining biological and clinical meaning by (3) exploration of brain regions and cell types implicated. These aims were all accomplished with the overarching goal of advancing biological understanding and identifying targets for (pharmacological and other) intervention and advancing precision medicine for PTSD.

## RESULTS

### GWAS and GWGAS of EHR Algorithmically Defined Case-Control PTSD

We performed genomewide association analysis (GWAS) of PTSD in American Veterans of European (EUR) and African (AFR) ancestry (initially separately, then meta-analyzed together) basing diagnosis on a validated VA EHR algorithm (Harrington et al., 2019) that had excellent discriminative ability for lifetime PTSD cases vs. controls as determined by chart review (0.90 sensitivity, 0.97 specificity, 0.87 positive predictive value, and 0.90 negative predictive value), and substantial agreement with gold-standard Clinician-Administered PTSD Scale (CAPS) interview (90.2% agreement and κ = 0.75 [95% CI: 0.62, 0.88]) (Radhakrishnan et al., 2019).

GWAS was carried out (on two tranches of data, distinguished by time when genotype results were available) on SNP dosages imputed from 1000 Genomes phase 3 using logistic regression in PLINK 2.0, separately by ancestry, adjusting for age, sex, and the first 10 principal components of ancestry. Meta-analysis by tranche (and later by ancestral group) was performed using METAL (Willer et al., 2010). The intent of our study was to conduct GWAS in combat exposed Veterans (Radhakrishnan et al., 2019), but combat exposure information was available for only a subset (51.2%) of the sample (**Table 1**), and GWAS of that subset yielded no genomewide significant (GWS) findings. However, genetic correlation [r_g_] between the categorical trait (i.e., diagnosis of) PTSD in those combat-exposed and in all subjects irrespective of combat exposure status was 0.969 ([se 0.049] p = 7.64×10^−89^), and therefore results for the latter larger, more informative, sample are presented here.

**Table 1.** Sample sizes and descriptive characteristics of participants.

|  | European Ancestry |  |  | African Ancestry |  |  |
| --- | --- | --- | --- | --- | --- | --- |
|  | Case | Control | All | Case | Control | All |
| N | 36301 | 178107 | 214408 | 11920 | 39116 | 51036 |
| Mean age (years) | 58.29 | 67.19 | 65.68 | 57.07 | 59.91 | 59.25 |
| Female | 6.6% | 5.2% | 5.6% | 12.0% | 10.8% | 11.1% |
| Combat exposure: |  |  |  |  |  |  |
| Exposed to combat | 40.0% | 25.0% | 27.5% | 23.9% | 9.4% | 12.8% |
| Not exposed to combat | 5.8% | 34.1% | 29.3% | 5.2% | 18.1% | 15.1% |
| Unknown exposure to combat | 54.2% | 40.9% | 43.1% | 70.9% | 72.5% | 72.1% |

The categorical trait (diagnosis) GWAS for the EUR sample included 36,301 algorithmically defined probable PTSD cases and 178,107 controls. Considering LD independent loci (r^2^ > 0.1), we identified three distinct GWS (p < 5×10^−8^) genomic risk loci (**Figure 1 [top]** and **Table 2**) on Chr11:28707675, rs10767744, (p=1.75×10^−10^) nearest to *METTL15*; on Chr7:70219946, rs137999048, (p=1.03×10^−8^) nearest to *AUTS2*; and on Chr7:1855531, rs7680, (p=4.17×10^−8^) nearest to *MAD1L1*, respectively.

**Table 2.** Genomewide significant (p < 5×10^−8^) findings with lead SNPs for EUR case-control GWAS (36,301 cases and 178,107 controls)

| LD Independent Lead SNP | Chr | Effect Allele (Frequency) | Odds Ratio (95% CI) | P-value | BP (GRCh37) | Nearest Gene |
| --- | --- | --- | --- | --- | --- | --- |
| rs10767744 | 11 | C (0.3911) | 0.945 [0.929-0.962] | 1.75E-10 | 28707675 | *METTL15* |
| rs137999048 | 7 | CCA (0.0473) | 1.123 [1.079-1.168] | 1.03E-08 | 70219947 | *AUTS2* |
| rs7680 | 7 | G (0.1411) | 0.931 [0.908-0.955] | 4.17E-08 | 1855531 | *MAD1L1* |

**Figure 1.**
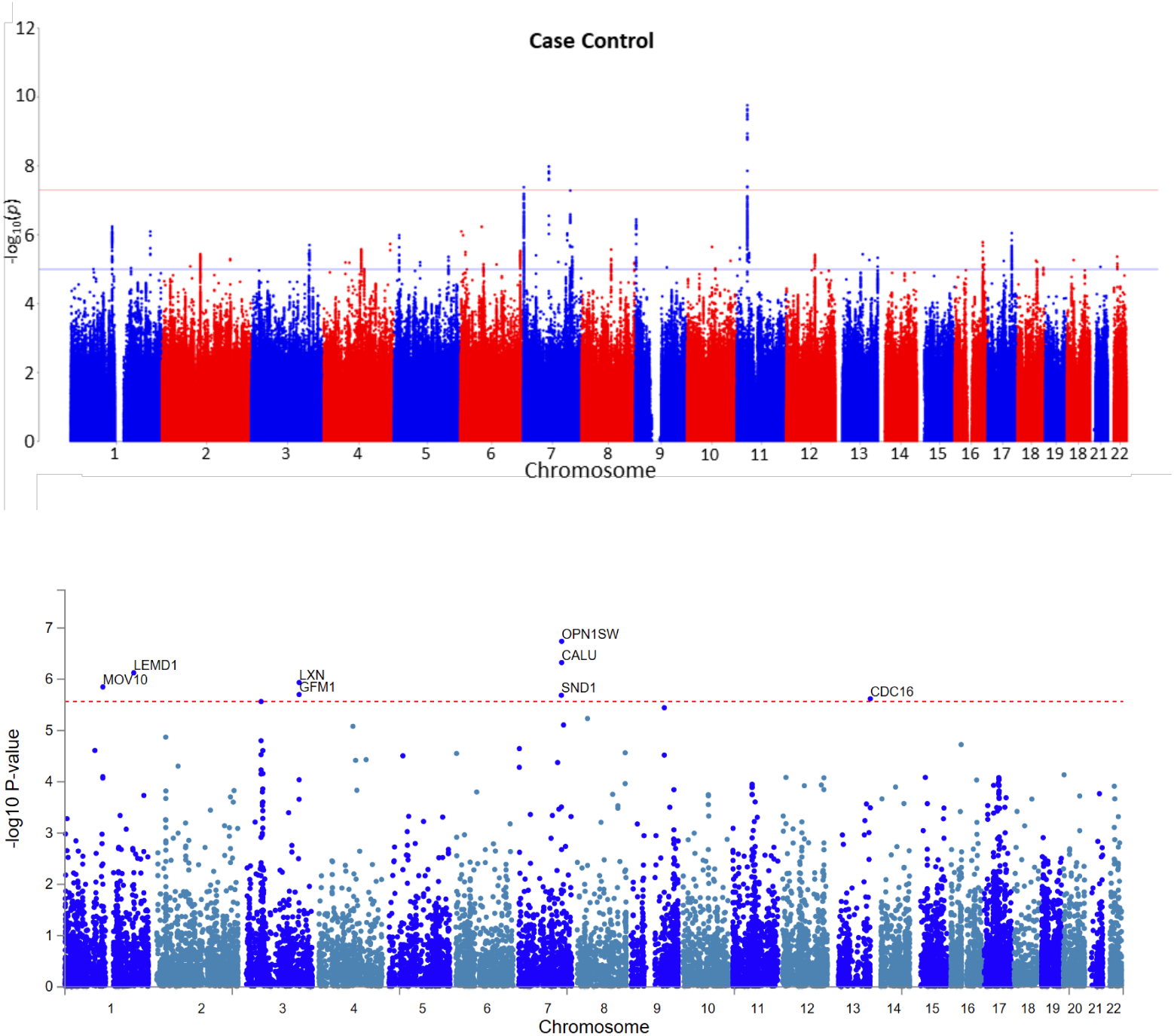
Top: Manhattan Plot of EUR Case-Control PTSD Phenotype GWAS Bottom: Manhattan Plot of EUR Case-Control PTSD Phenotype GWGAS

We also conducted genomewide gene-set-based association tests (GWGAS) using MAGMA (de Leeuw et al., 2015), which identified 8 additional genes (**Table 3** and **Figure 1 [bottom]**) significantly associated with PTSD diagnosis.

**Table 3.** Genomewide significant (p = 0.05/17,927 = 2.79×10^−6^) findings for EUR PTSD case-control GWGAS.

| GENE | CHR | BP START | BP STOP | SNPS (N) | Z SCORE | P-value |
| --- | --- | --- | --- | --- | --- | --- |
| *OPN1SW* | 7 | 128412545 | 128415844 | 13 | 5.086 | 1.83E-07 |
| *CALU* | 7 | 128379346 | 128411861 | 86 | 4.9023 | 4.74E-07 |
| *LEMD1* | 1 | 205350506 | 205425082 | 270 | 4.8115 | 7.49E-07 |
| *LXN* | 3 | 158363611 | 158390482 | 102 | 4.7231 | 1.16E-06 |
| *MOV10* | 1 | 113215763 | 113243368 | 63 | 4.6828 | 1.41E-06 |
| *GFM1* | 3 | 158362067 | 158410364 | 187 | 4.6128 | 1.99E-06 |
| *SND1* | 7 | 127292234 | 127732661 | 787 | 4.6052 | 2.06E-06 |
| *CDC16* | 13 | 115000362 | 115038198 | 108 | 4.5731 | 2.40E-06 |

The GWAS for the AFR sample included 11,920 algorithmically-defined probable PTSD cases and 39,116 controls. We identified two distinct GWS loci, one on Chr3:1259951, rs4684090 (p=3.59×10^−8^) intronic to *CNTN6* and one on Chr20:6724577, rs112149412 (p=3.18×10^−8^) near *BMP2*. GWGAS using MAGMA identified two additional genes (*MAPK13* on Chr6 and *CDC14B* on Chr9).

GWAS for the 48,221 cases and 217,223 controls in the trans-ancestral analysis (i.e., meta-analysis of EUR and AFR samples) (data not shown) identified as GWS SNPs in two of the same genes found GWS in the EUR GWAS: a different lead SNP on Chr7: 1959634 (rs137944087, a common indel/deletion) in moderate LD with the variant identified in the EUR sample (r^2^=0.38, D’=0.78), and a different lead SNP on Chr11: 28678870 (rs10767739) in LD with the variant identified in the EUR sample (r^2^=0.54, D’=0.84).

### GWAS of PTSD Symptom Sub-phenotypes and Total Symptoms

The MVP surveys included the PTSD Checklist for DSM-IV (PCL), a widely used 17-item self-report measure of past-month PTSD symptoms covering the 3 DSM-IV diagnosis components – re-experiencing, avoidance, and hyperarousal – as well as a total symptom severity score as the sum of those 3 sub-phenotypes (Blanchard et al., 1996). GWAS with these phenotypes in the EUR sample (N = 186,689 individuals) using linear regression revealed multiple GWS loci including some that were associated with total symptom score as well as multiple symptom sub-domains (e.g., *MAD1L1*; *TCF4; TSNARE1*), and others that were more strongly associated with certain symptom sub-domains (e.g., *CAMKV* to re-experiencing; *SOX6* to hyperarousal) (**Table 4**). The overlap in risk loci for the 3 sub-phenotypes and their total is shown in **Figure 2** (and as stacked Manhattan plots in **Figure 3**).

**Table 4.**
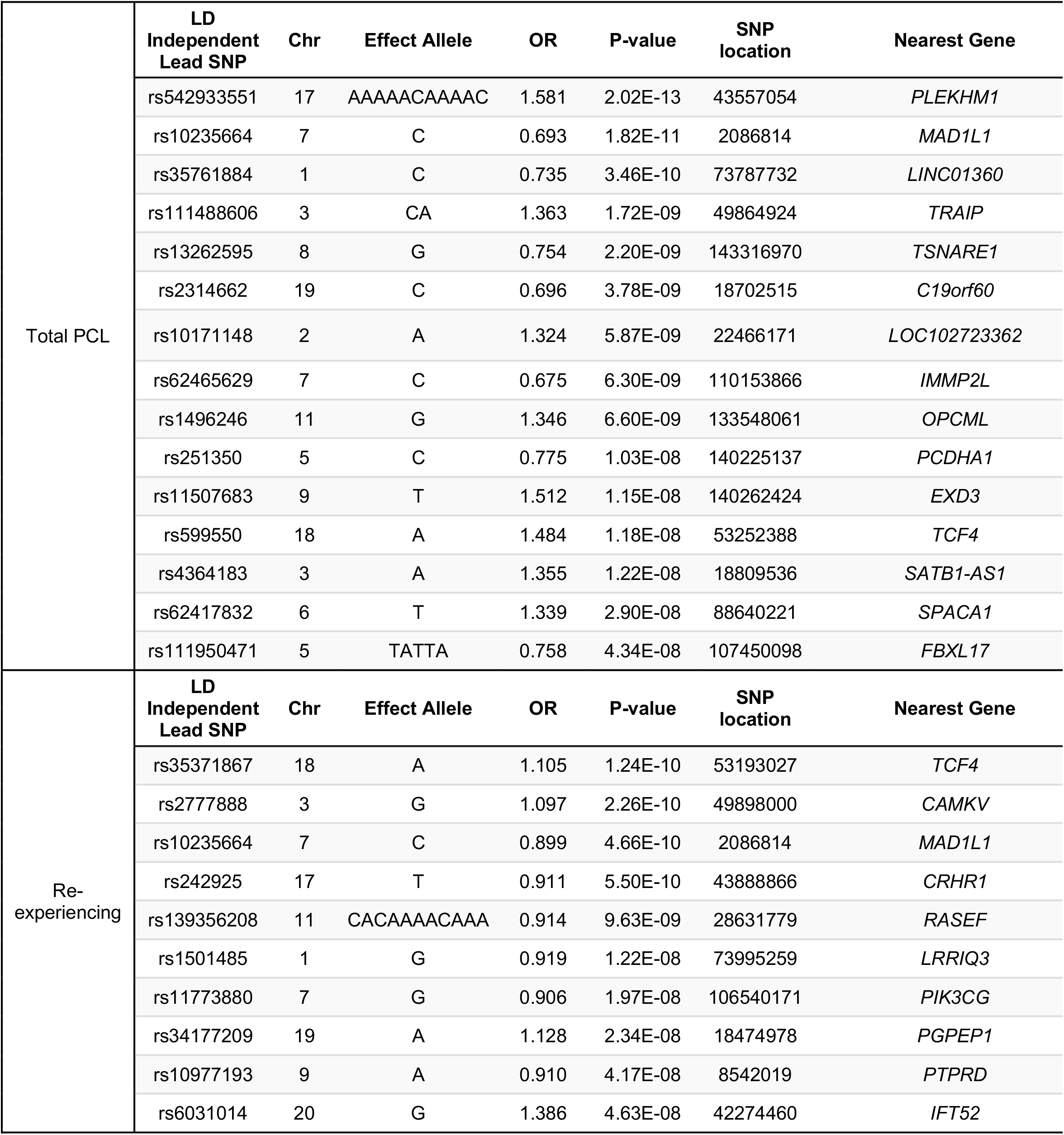
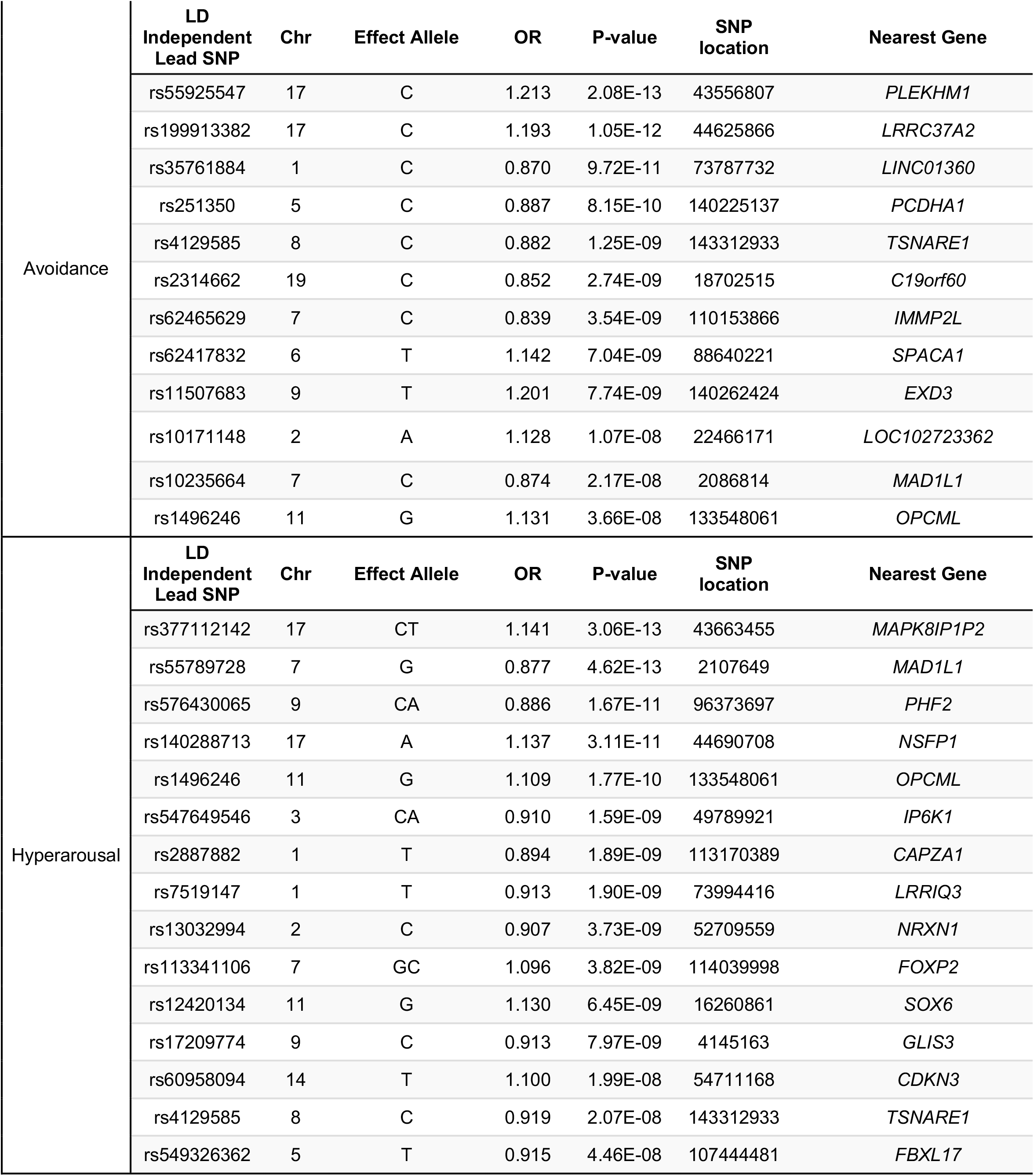
Genomewide significant (p < 5×10^−8^) findings with lead SNPs for EUR PCL Total and Sub-Phenotype GWASs (N = 186,689 individuals)

**Figure 2.**
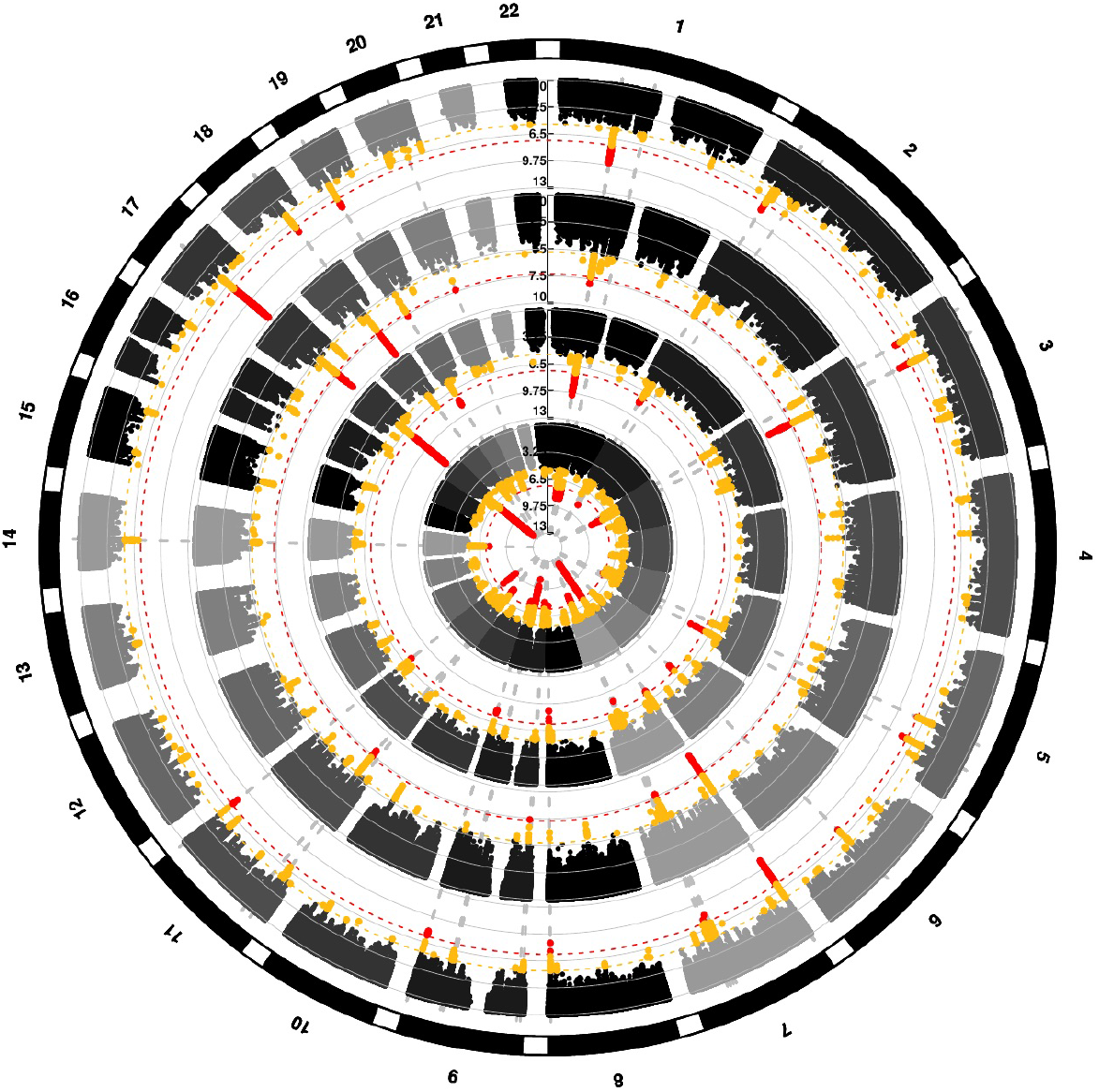
Circle Manhattan Plot for PCL phenotypes in European Americans. The results, from outermost to innermost circle, are Total PCL, Re-experiencing, Avoidance, and Hyperarousal. Red dots indicate genome-wide significant findings (p<5×10^−8^) and yellow dots indicate suggestive findings (p<5×10^−6^). The numbers around the circle indicate the chromosome. Vertical dashed grey lines are drawn through genome-wide significant findings to indicate overlap between analyses.

**Figure 3.**
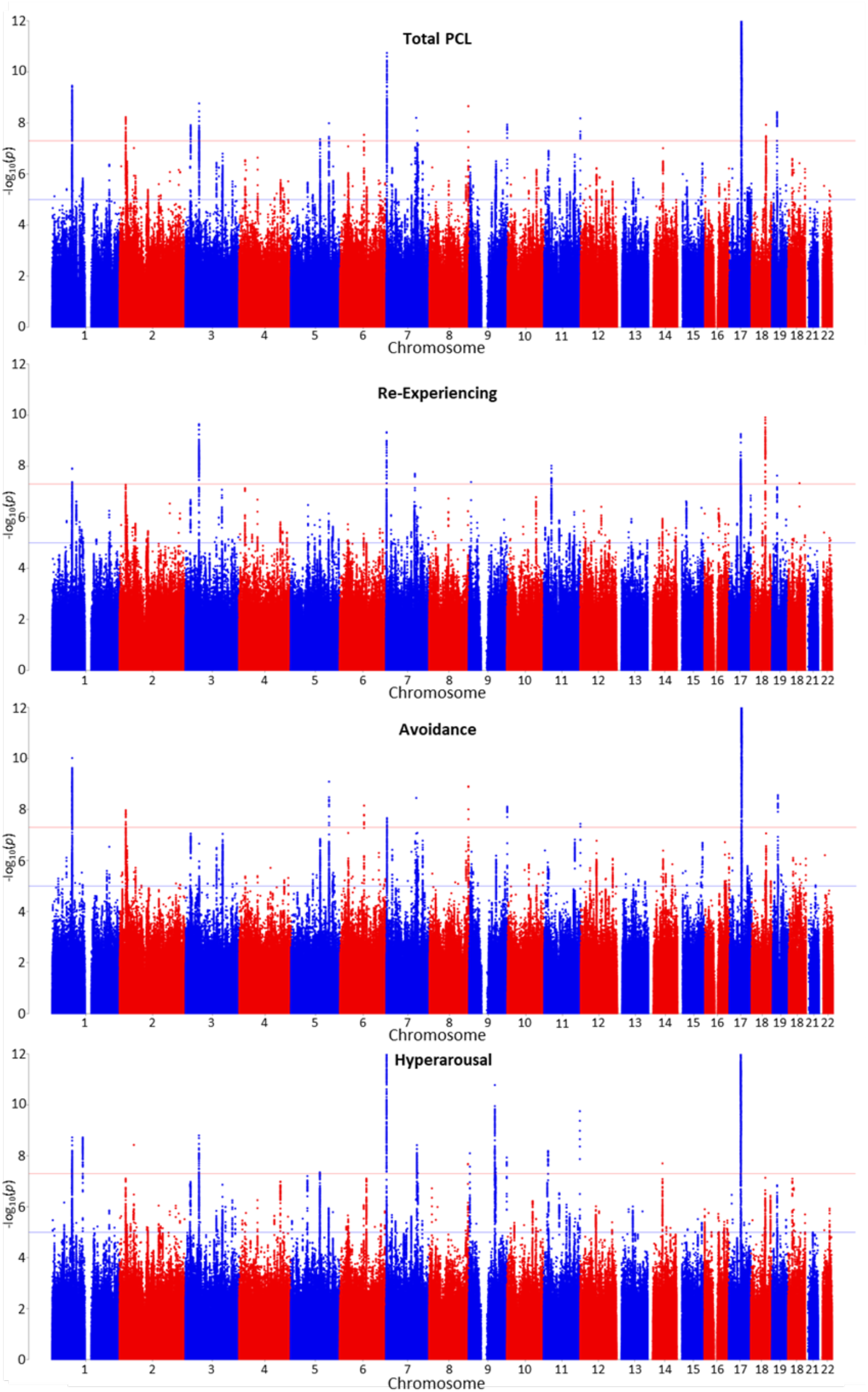
Stacked Manhattan Plots for all 4 traits (from top to bottom: total symptoms, re-experiencing, avoidance and hyperarousal) in the EUR sample

MAGMA identified 77 genes that were significant (p = 0.05/15659 = 3.19×10^−6^) in the gene-based analysis of PCL-total score in EUR (**Supplemental Table 1**). Virtually all these genes also had SNPs associated with PCL-total and/or sub-phenotype scores in the EUR GWAS (e.g., *MAD1L1*, *TCF4*, *PLEKHM1*, *CRHR1*) although several new genes were also implicated (e.g., *WNT3*, which is part of the LD block on Chr17 that also includes *CRHR1*).

### B. Heritability of PTSD Phenotypes and Partitioned Heritability

SNP-chip heritability (on the liability scale, assuming 10% prevalence of PTSD in US Veterans) for MVP-PTSD (algorithmic case-control) in EUR using LDSC is 0.064 (se 0.0055). SNP-chip heritability estimates (on the observed scale) in EUR for the 4 quantitative traits were not significantly different from one another: PCL-total (0.092 [se 0.0052]), PCL-reexperiencing (0.093 [se 0.005]), PCL-avoidance (0.093 [se 0.0055]), and PCL-hyperarousal (0.101 [se 0.0058]). Heritability was significantly stronger for the PCL-total (z-score = 17.73) than for the algorithmic case control phenotype (z-score = 11.62) (p=0.0002). Subsequent post-GWAS analyses were therefore conducted, unless otherwise specified, on the most genetically informative phenotype, i.e., the PCL-total quantitative trait.

Partitioning heritability of PCL-total in EUR revealed 1.32- fold to 1.38-fold enrichment of SNPs associated with three GTEx cortical tissue types: brain cortex, brain frontal cortex (BA9), and brain anterior cingulate cortex (BA24) (FDR p < 0.05); intronic regions had 1.27-fold enrichment (p=2.78×10^−4^). Cell-type analyses support heritability enrichment of the frontal cortex (BA9) gene sets (tau-c=4.64×10^−9^, p=0.002) and frontal cortex (BA9) and anterior cingulate cortex (BA24) gene expression profiles (BA9 tau-c=5.30×10^−9^, p=8.41×10^−4^; BA24 tau-c=6.28×10^−9^, p=2.26×10^−4^), above that of all other genomic annotations. (Data not shown.)

### C1. Phenotypic and Genetic Correlations Across PTSD Phenotypes

**Table 5** shows the phenotypic (above the diagonal) and genetic (below the diagonal) correlations in EUR between the algorithmic case-control diagnosis, and each of the 4 continuous PTSD measures (reexperiencing, avoidance, hyperarousal, and their total). As shown, even when phenotypic correlations were modest, genotypic correlations were consistently high (> 0.9).

**Table 5.**
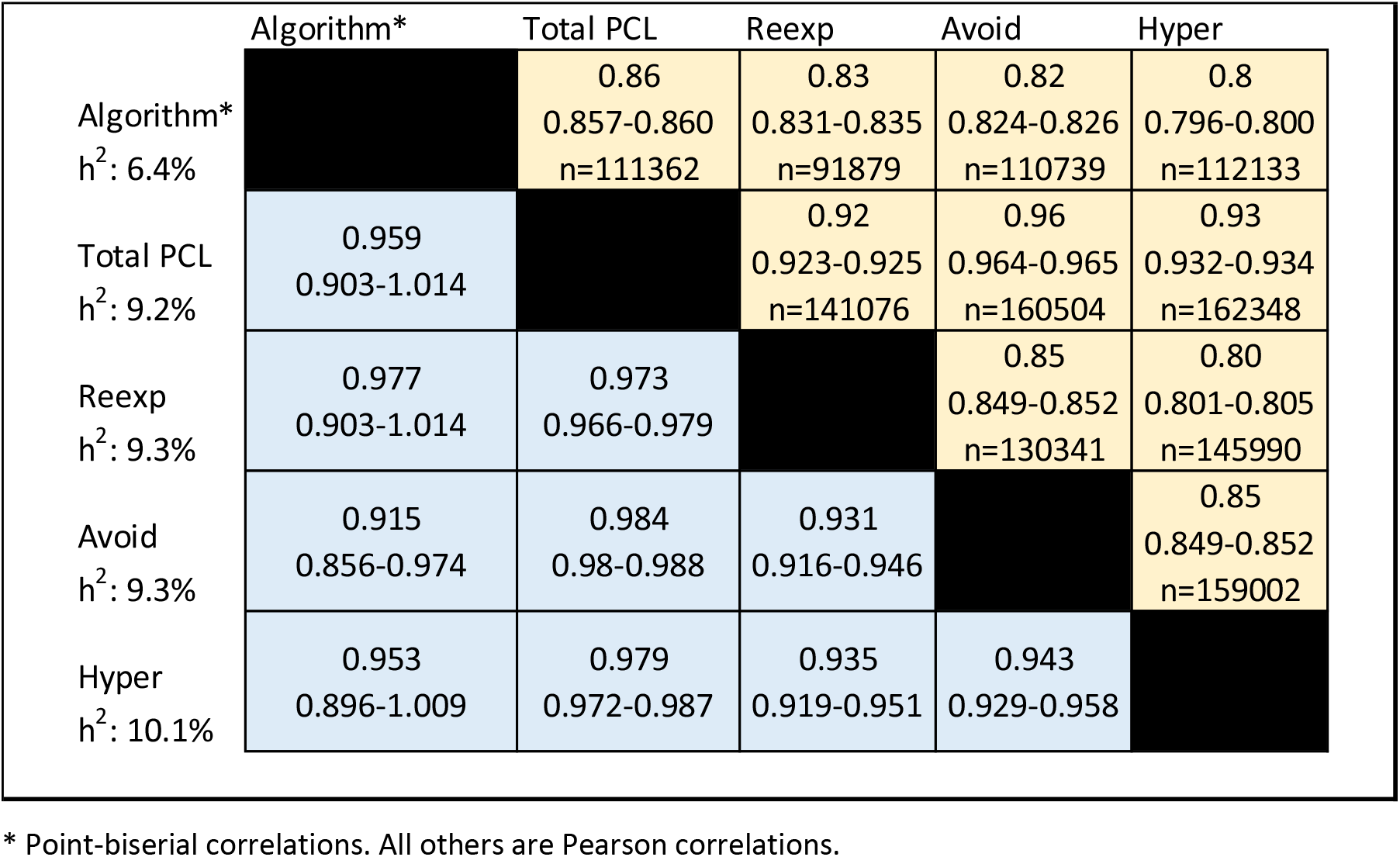
Phenotypic (above diagonal) and genetic (below the diagonal) correlations between algorithmic case-control diagnosis, PCL-total, and PCL-sub-scores. Shown are point estimates for correlations, 95% CIs, and n (sample size)

### C2. Genetic Correlation Using LD Score Regression in External Datasets

In the EUR sample, we estimated genetic correlations between PCL-total score and health-related traits available in LDHub. The many significant genetic correlations included: depressive symptoms (r_g_ = 0.741, p = 2.26×10^−67^), neuroticism (r_g_ = 0.637, p = 2.07×10^−47^), intelligence (r_g_ = −0.460, p = 5.26×10^−34^), subjective well-being (r_g_ = −0.393, p = 4.90×10^−21^), and insomnia (r_g_ = 0.489, p = 9.15×10^−20^) (**Figure 4** and **Supplemental Table 2a**).

**Figure 4.**
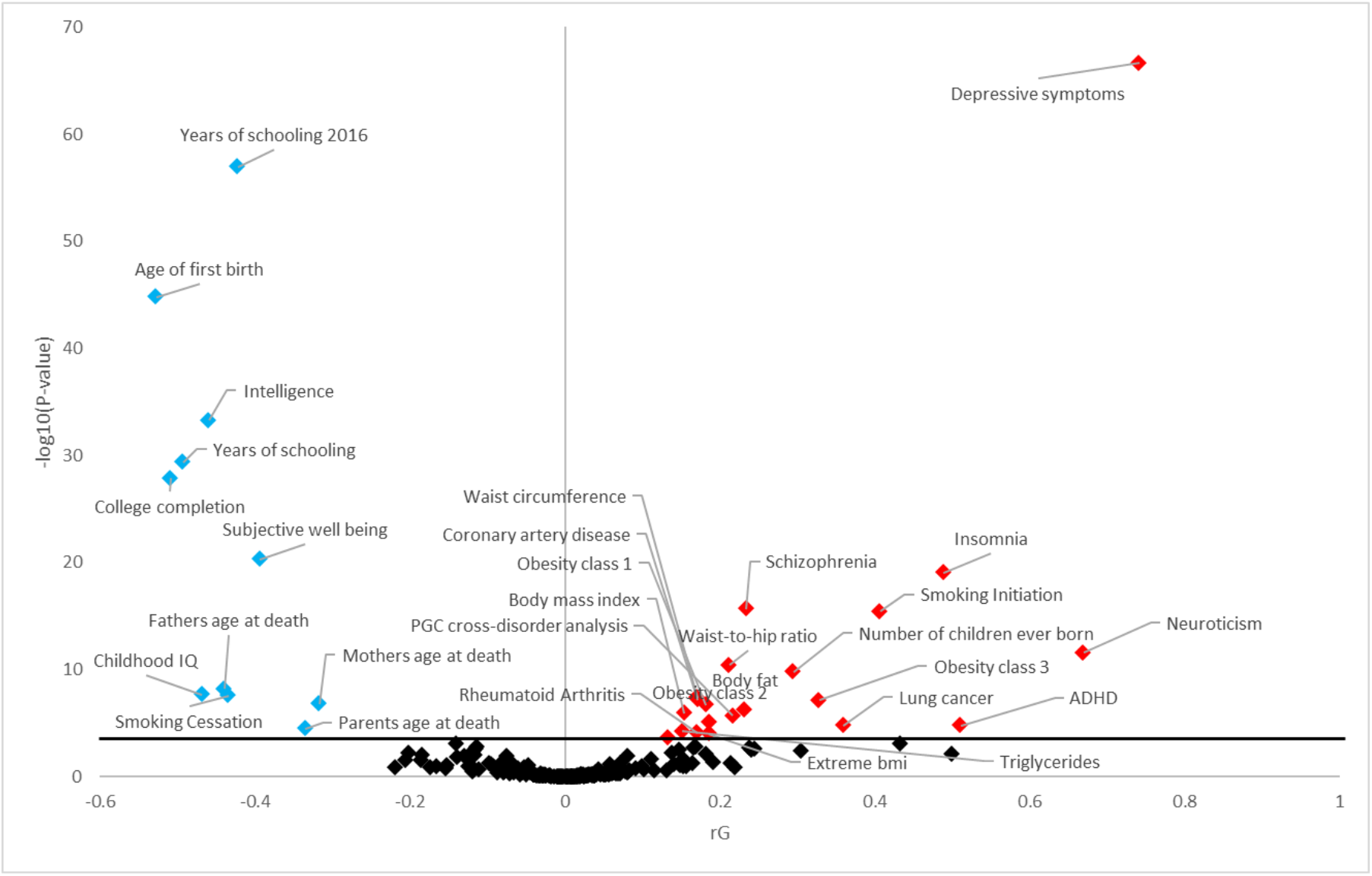
LDSC analyses showing traits from LDHub with the most significant genetic correlations (r_g_ on the x-axis and −log10 p-value [Bonferroni-corrected] on the y-axis) with PCL-total in EUR. Blue diamonds indicate statistically significant negative r_g_; red diamonds indicate statistically significant positive r_g_; black diamonds indicate non-significant r_g_.

### C3. Replication of GWAS Findings

We compared our top SNP associations from the PTSD case-control and PCL-total results against the largest available PTSD external dataset, PGC-PTSD 2.0 (Nievergelt et al., 2018). For our case-control phenotype, we used the lead SNPs in MVP, and for our continuous PTSD symptom scores (PCL) we considered independent GWS variants (r^2^<0.1).

For the EUR case-control phenotype, there was nominal replication for 1 of 3 SNPs: for rs7680*A nearest to *MAD1L1*, with a beta of −0.0712 (se 0.013, p=4.17×10^−8^) in MVP and a beta of −0.0639 (se 0.0215, p=0.00312) in PGC-PTSD 2.0. For the EUR PCL-total symptom scores, there were 6 of 15 possible nominal replications (including, notably, in *MAD1L1*, *TSNARE*, and *EXD3*). Details are provided in **Supplemental Table 3**.

We also applied a PRS in EUR for the algorithmic case-control phenotype and the PCL-total quantitative trait. Both PRSs strongly and significantly predicted into PGC-PTSD 2.0 with the MVP case-control PRS explaining approximately 0.4% of the variance (p=2.4×10^−74^) in the PGC-PTSD 2.0 phenotype at p-value thresholds (p_T_) > 0.05, with the MVP PCL-total PRS explaining 0.7%-0.8% of the variance (p=2.2×10^−134^) in the PGC-PTSD 2.0 phenotype at p_T_ > 0.05 (**Figure 5**).

**Figure 5.**
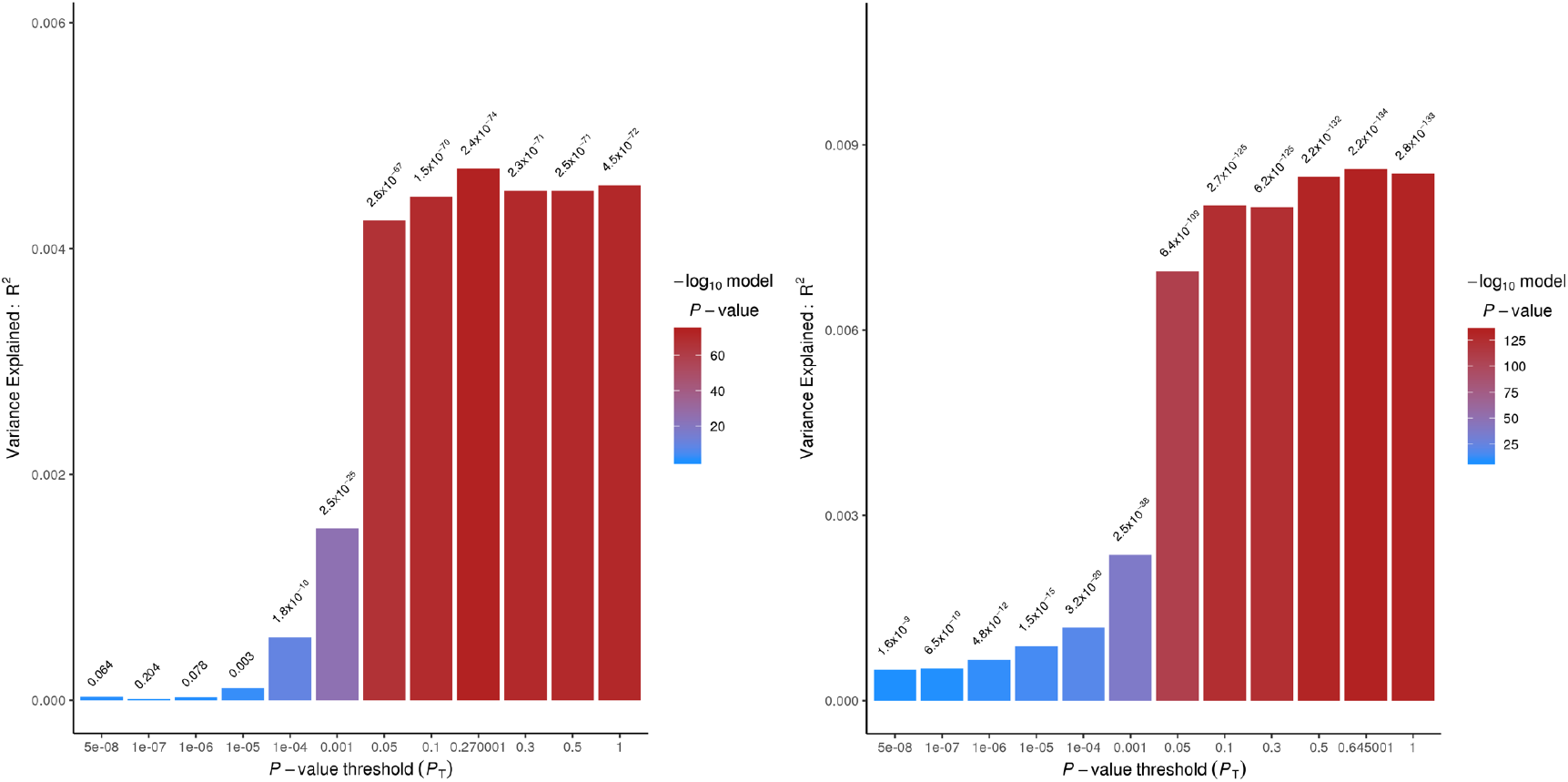
Polygenic risk score (PRS) from MVP EUR case-control (left) and EUR PCL-total (right) applied to PGC-PTSD 2.0 case-control phenotype with varying P-value thresholds (P_T_) on x-axis and explained (R^2^) on y-axis.

### D. Enrichment in Biological Tissues and Pathways Using MAGMA and PrediXcan

Results from MAGMA, using the PCL-total EUR results revealed significant enrichment according to GTEx V7 in two (cerebellum and cerebellar hemispheres) of 57 tissues.

PrediXcan-S (Barbeira et al., 2018) was used to correlate tissue-specific expression determined by association with reference transcriptome datasets with the PCL-total quantitative trait results. Significant *negative* correlation with predicted expression of the protein product of the pseudogene *LRRC37A4P* (*aka LRRC37* and *LRRC37A4*) in brain (amygdala, substantia nigra, putamen/basal ganglia, multiple cortical regions including anterior cingulate), adrenal gland, and whole blood was observed. Also noted was significant *positive* correlation with predicted expression of *CRHR1* (notably CRHR1-IT1-CRHR1 readthrough protein) in brain (amygdala, hippocampus), adrenal gland and whole blood. (Data not shown.)

### E. Postmortem Brain Findings in Subgenual Cingulate Cortex

Following up on the PrediXcan-S results for *CRHR1*, postmortem brain tissue from subgenual prefrontal cortex (BA25) from subjects with PTSD matched to controls were assayed for gene expression changes by quantitative real-time PCR. Normalized log-corrected Ct values for *CRHR1* were used to calculate fold change. This analysis revealed a significant 1.3-fold increase in *CRHR1* expression (**Figure 6**) (Bonferroni-corrected P< 0.001, n=22, error bars indicate ± SEM). There were no significant differences in gene expression between subjects with PTSD who were medicated and nonmedicated at time of death.

**Figure 6.**
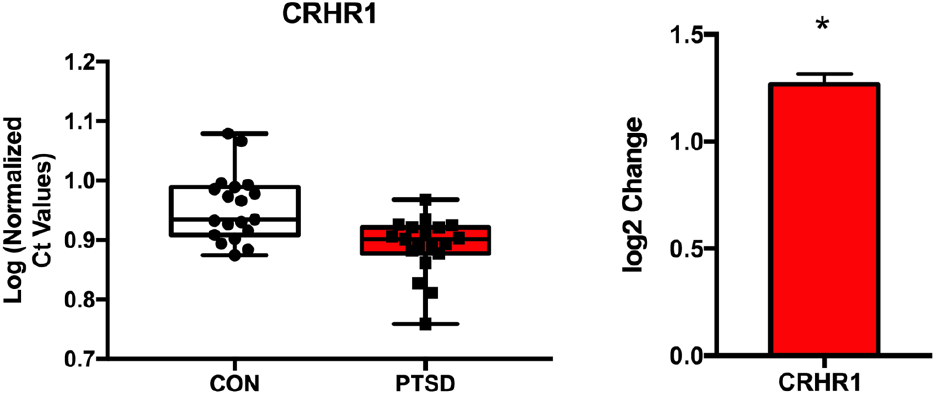
Gene expression analysis of target genes in postmortem human BA25. A) Box plots of normalized Ct values calculated for CRHR1 in controls (white) and PTSD (red) subjects. B.) Log 2-fold change of CRHR1 in human subgenual PFC (Bonferroni corrected, *P*<0.001, n=22). Error bars indicate ± SEM.

### F. Genomic Relationship of PTSD to Major Depression and Other Major Mental Disorders

We used mtCOJO (Zhu et al., 2018) to address the genetic relationship between PTSD and other major mental disorders. By applying the most recent PGC GWAS results for 8 mental health traits — autism spectrum disorder, major depression, anorexia nervosa, anxiety (case-control), alcohol dependence, schizophrenia, bipolar disorder, attention deficit hyperactivity disorder (Duncan et al., 2017; Grove et al., 2019; Howard et al., 2019; Martin et al., 2018; Otowa et al., 2016; Schizophrenia Working Group of the Psychiatric Genomics, 2014; Stahl et al., 2019; Walters et al., 2018) — and all the aforementioned traits to our PCL-total quantitative phenotype in EUR, we were able to parse out the genetic signal attributable to PTSD alone. PCL-total remained highly genetically correlated with the unconditioned GWAS when conditioned on genetically correlated psychiatric disorders independently and simultaneously (**Figure 7**).

**Figure 7.**
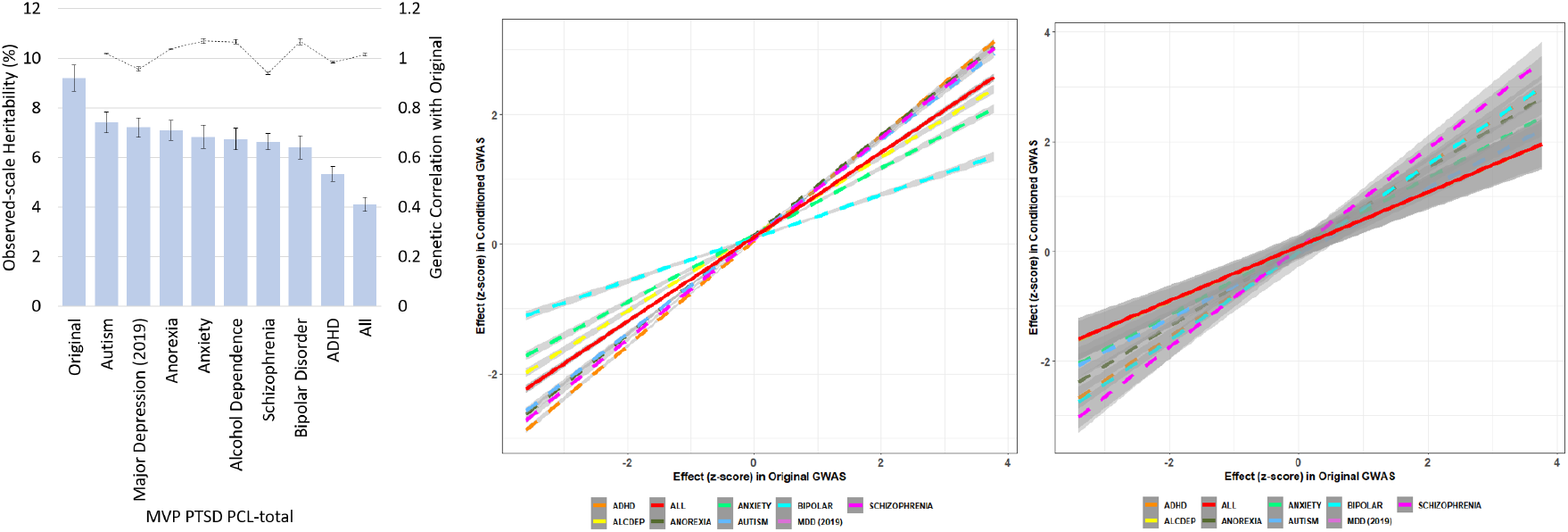
First figure shows observed scale h^2^ and r_g_ relative to original PCL-total. Second figure shows linear relationship between GO term enrichment in original PCL-total and conditioned PCL-total. Third figure shows linear relationship between GTEx tissue enrichment in original PCL-total and conditioned PCL-total.

Conditioning with all 8 mental disorder traits significantly reduced the observed-scale heritability of PCL-total by 5.10% (PCL-total original h^2^ = 9.21%, p=1.39×10^−67^; PCL-total conditioned h^2^ = 4.11%, p=2.61×10^−52^) relative to the unconditioned GWAS (p_difference_=1.52×10^−13^), but this reduction in heritability did not significantly alter associations with biological pathways or tissues associated with genetic risk for PTSD (**Figure 7**).

### G. PheWAS

A phenome-wide association study (PheWAS) within the VA-MVP EHR sample (N=381,609) of the top GWS SNPs from the PCL-total quantitative trait GWAS revealed several directionally consistent (i.e., effect allele associated with increased risk for the disorder and for increased PTSD symptoms) and significant (p<3.75×10^−5^) associations with a group of mental disorders (e.g., “anxiety, phobic and dissociative disorders” (p=8.74×10^−6^) in the case of rs11507683 on Chr9 near *EXD3;* and with mood disorders (p=5.3×10^−7^) and tobacco use disorder (p=1.15×10^−6^) in the case of rs35761884 on Chr1 in *LINC01360*). Also noted were directionally consistent and significant associations with several physical disorders, including rs542933551 (on Chr17 in *PLEKHM1*) with hypothyroidism (p=4.91×10^−14^) and kidney stones (p=2.12×10^−8^); and of rs111488606 (near *TRAIP* on Chr3) with chronic airway obstruction (p=1.47×10^−5^). (Data not shown.)

### H. Drug Repositioning Analysis

We selected the GWS genes from the gene-based analysis (conducted using MAGMA) of PCL-total, filtered the list by those genes that were also GWS in the GWAS for this trait (including *CRHR1*, which was part of a large LD block in Chr17 and also was prominently implicated in the PrediXcan-S analyses, noted above), and added *METTL15* and *AUTS2* which were GWS in the case-control analysis. We then imported this list of 10 genes (*MAD1L1*, *METTL15*, *AUTS2*, *TSNARE1*, *EXD3*, *PLEKHM1*, *TCF4*, *TRAIP*, *C19orf60*, *CRHR1*) into the Drug Gene Interaction Database v3.0 (dgidb.genome.wustl.edu) (Cotto et al., 2018) to determine if there were interactions with available drug treatments that might indicate potential novel drug strategies for PTSD. Drug repositioning analysis was also carried out in the Connectivity Map (CMap) database (https://www.broadinstitute.org/connectivity-map-cmap) for the same set of 10 genes (Subramanian et al., 2017). *CRHR1* was identified in both databases as being a potential drug target with experimental medications available. Given the positive association between PTSD symptoms and imputed *CRHR1* expression in brain, in concert with the direct observation of upregulated *CRHR1* expression in postmortem PTSD brain, a *CRHR1* antagonist would be hypothesized to be potentially therapeutic. Also identified was *TCF4* in association with drugs such as darinaparsin (an apoptosis stimulant under investigation to treat some cancers). CMap also indicated strong functional overlap (based on knockdown expression profiles) of *TCF4* with *PLXNA1* (CMap expression similarity score = 98) which has as a druggable target the acetycholine M2 receptor antagonist otenzepad. Another gene, *PLEKHM1*, which was among the top 3 genes (p = 1.39×10^−11^) in the GWGAS for PCL-total, was considered by CMap as highly likely to share biological effects with several classes of drugs, including dopamine receptor antagonists, acetylcholine receptor antagonists, and angiotensin receptor antagonists.

## DISCUSSION

The past decade has seen a proliferation in the use and usefulness of GWAS, with the prediction that continued sample size growth will result in even richer findings (Visscher et al., 2017). The field of psychiatric genomics has capitalized on the uses of GWAS, with substantial gains made in the understanding of serious mental disorders such as schizophrenia, major depression, and bipolar disorder (Smoller, 2019; Sullivan et al., 2018). The utility and interpretability of the results depend on the data source, including the specificity of available phenotype information and the homogeneity of the sample; or, if it is heterogenous, whether it is heterogenous in known ways. Notably, PTSD has lagged behind these other disorders in assembling adequately powered studies, though recent advances are noteworthy (Duncan et al., 2018; Gelernter et al., 2019; Nievergelt et al., 2018; Stein et al., 2016). We present here the largest single-source case-control GWAS of PTSD to date, and we augment its informativeness with the GWAS of several other self-report PTSD phenotypes, including a quantitative trait corresponding to symptom severity, which proved more genetically informative than the case-control analysis. We examined whether traits that correspond to the main PTSD symptom groups employed in clinical diagnosis are related genetically, and the extent to which they are related phenotypically in this sample, the largest-ever studied for this purpose. This study is the first to compare and contrast genetic risk for PTSD from various phenotypic perspectives and, to the best of our knowledge, the first to robustly replicate a polygenic signal in an independent case-control sample and, importantly, to replicate (nominally) in several specific, newly-identified variants (e.g., in *MAD1L1*) in external datasets. These analyses revealed several genomewide significant (GWS) associations with PTSD visible at the case-control level, and numerous GWS associations with various dimensions of symptom severity (which included more subjects and more information per subject, and therefore were better powered than the analysis based on diagnosis). When combined with imputed expression results and enrichment analyses, these results help to illuminate the neurobiology of PTSD and, importantly, begin to uncover new avenues for therapeutic development.

Derived from the uniquely informative US Veterans Affairs MVP (Gaziano et al., 2016), our case-control sample was phenotyped using a validated algorithm applied to EHR data with exemplary diagnostic properties (Harrington et al., 2019; Radhakrishnan et al., 2019), enabling us to analyze data from over 250,000 genotyped veterans (including nearly 50,000 PTSD cases). Moreover, we were able to examine simultaneously genomewide associations with several other PTSD phenotypes reflecting various aspects of symptom type and severity, respectively. These data highlight the tremendous scientific value of EHR-based biobanks and automated phenotyping for genomic research (Chen et al., 2018; Smoller, 2018), and even more so, the value of collecting relevant self-report data such as the PCL. Biobanks have specific subject and ascertainment characteristics; the nature of the MVP makes it uniquely suitable for study of PTSD, a disorder that occurs at increased rates in military veterans.

This is the first study to directly compare heritability of binary (diagnostic) and continuous (symptom-based, including symptom subsets) phenotypes for PTSD. We found that both were significantly heritable, though the continuous (total symptom) trait was the most heritable, and therefore the most informative with regard to biological inference. Importantly, partitioned heritability analyses of that trait indicated preferential expression of SNPs in frontal (BA9) and anterior cingulate cortex (BA24), consistent with prevailing neural circuit theories of PTSD pathophysiology (Shalev et al., 2017) that emphasize function of these regions and their connections with limbic cortex in the regulation of emotion and extinction of fear memories (Dunsmoor et al., 2015; Phelps and Hofmann, 2019).

Several genes were repeatedly implicated across the various conceptualizations of the PTSD phenotype. *MAD1L1* (“mitotic arrest deficient 1 like 1”) was GWS associated with PTSD in the algorithmic case-control comparison, with all three symptom severity dimensions (hyperarousal, reexperiencing [previously seen and now remaining GWS in this larger sample [(Gelernter et al., 2019)] and avoidance) and also with total symptom severity in the quantitative trait analyses including the gene-based analysis. *MAD1L1*, widely expressed in all tissues and thought to play a role in cell cycle control, has emerged as being GWS associated with at least two other major mental disorders, schizophrenia (Schizophrenia Working Group of the Psychiatric Genomics, 2014) and bipolar disorder (Stahl et al., 2019) — both of which traits were excluded among participants in this study but have strong genetic correlations with PTSD in MVP and other cohorts (Nievergelt et al., 2018). These observations suggest that *MAD1L1* may be a general risk factor for psychopathology, possibly contributing to the *p factor* thought to underlie many serious mental disorders (Selzam et al., 2018). Alternatively, it could contribute to some of the symptoms or clusters of symptoms that these disorders can have in common. These hypotheses should be directly tested in future studies that focus on cross-disorder psychopathology.

Several other genes were discovered as being associated with PTSD and, remarkably, replicated in the largest available (but still much smaller than our study sample) PTSD-informative dataset, the updated PGC-PTSD GWAS (Nievergelt et al., 2018). Included among these were *TSNARE1* (T-SNARE Domain Containing 1) and *EXD3* (Exonuclease 3’-5’ Domain Containing 3)*. TSNARE*, involved in intracellular protein transport, has been associated with risk-taking (Karlsson Linner et al., 2019), which may predispose to PTSD through increasing the likelihood of exposure to traumatic events; interestingly, twin studies suggest that risk for exposure to traumatic events is partially heritable (Stein et al., 2002). *EXD3*, involved in nucleic acid binding and widely expressed throughout the body, has been associated with mathematical (Lee et al., 2018) and other cognitive abilities, which have been found in our study and others to be genetically correlated with PTSD and mediated by socioeconomic status (Polimanti et al., 2019). It remains to be determined to what extent these associations and their potential mediation processes reflect pleiotropic effects of *EXD3* or other genes. But, for the first time in PTSD genetics research, we have discovered and replicated gene candidates through unbiased searches that can now be further examined in relation to their putative biological relationships to PTSD and other stress- and anxiety-related conditions.

Our prior analysis of intrusive reexperiencing symptoms in MVP (Gelernter et al., 2019) had implicated *CRHR1*, and this gene was once again implicated in this expanded sample both through its association with the same quantitative trait in the GWAS (Table 3), and also through its strong association with overall PTSD symptoms (PCL-total) in the gene-wise analysis (Supplemental Table 1). *CRHR1* is in a large LD block on Chr17, making it difficult to discern its association with PTSD apart from other genes in that LD block. We now provide additional biological evidence that *CRHR1* may be causally related to PTSD. PrediXcan analyses pointed to heightened expression of *CRHR1* in several brain regions (amygdala and hippocampus) often implicated as structurally or functionally abnormal in PTSD (Shalev et al., 2017). And, importantly, our postmortem brain data directly demonstrate increased expression of *CRHR1* in individuals with PTSD. Limitations of these data include the availability of postmortem tissue only in BA25, and many other caveats that go with postmortem brain analysis (e.g., confounds possibly caused by age, other comorbid conditions, and medications). These results must be extended to other brain regions (e.g., ventromedial prefrontal cortex, shown to be integral to fear learning and extinction (Dunsmoor et al., 2019), processes hypothesized to be central to PTSD onset and recovery, respectively (Maddox et al., 2019; Shalev et al., 2017)) and replicated in additional samples. But the totality of the results now associating *CRHR1* with PTSD severity across GWAS, GWGAS, imputed expression, and postmortem analyses, in concert with the strong preclinical and clinical priors for involvement of CRH in stress-related disorders (Chrousos and Zoumakis, 2017), position *CRHR1* antagonists as strong therapeutic candidates for PTSD and related conditions. Whereas a placebo-controlled trial of a CRHR1 antagonist in 128 women with PTSD produced unimpressive results (Dunlop et al., 2017), our findings (albeit predominantly in men) suggest that there are potential unfulfilled opportunities with *CRHR1* antagonists for PTSD that should be further explored. When such studies are performed, consideration should be given to looking at individual variation in *CRHR1*, including epigenetic variation (Pape et al., 2018), as a source of differential antagonist efficacy, in keeping with the march toward precision psychiatry (Stein and Smoller, 2018).

Our observation of the high r_g_ among the PCL sub-domains warrants additional reflection. When traits have high phenotypic correlations (r_p_) it is virtually always the case that they have high r_g_ (Sodini et al., 2018). Therefore, it may be somewhat unreasonable to expect that different levels of observation or measurement of traits that are highly phenotypically correlated will yield new insights into genetic risk (and, hence, neurobiology). The high r_g_ between PTSD symptom subdomains, which do not include overlapping items, supports the coherence of PTSD as a diagnostic construct from a biological perspective. Nevertheless, we did see evidence of certain SNPs being associated with risk for some PTSD sub-domains but not others, suggesting that there is further merit to looking at these sub-phenotypes as a tool for understanding disorder biology. This approach is consistent with expectations for sets of symptoms that, although they appear disjointed, tend to occur together and can be used to arrive at diagnoses that have predictive value. It is also possible that SNPs that are subdomain-specific in this dataset may not remain so as sample sizes increase and new findings emerge for each trait; but we would still expect that different, or differential, patterns for each trait will be maintained. It will be important to go beyond documentation of genetic correlation between PTSD and other traits and do the work needed to determine which of these relationships are causal and in which direction. Methodologies for such analyses have been developed (e.g., Mendelian Randomization (Emdin et al., 2017) and latent causal variable (LCV) modeling (O’Connor and Price, 2018)), and we intend to apply these techniques to the large MVP dataset to further elucidate causal risk factors for PTSD and related conditions.

PTSD is frequently associated with other mental health problems such as major depressive disorder (Stander et al., 2014), nicotine (Kearns et al., 2018) and alcohol (Vujanovic et al., 2019) abuse, and suicidality (Nock et al., 2009; Ramsawh et al., 2014) and with other adverse health sequelae such as obesity (Kubzansky et al., 2014) cardiovascular disease (Koenen et al., 2016), dementia (Cohen et al., 2013), type 2 diabetes (Roberts et al., 2015) and other immune-related disorders such as rheumatoid arthritis (O’Donovan et al., 2014) and hypothyroidism (Jung et al., 2018). Our PheWAS yielded a significant association between hypothyroidism in MVP with rs542933551 (in *PLEKHM1*). Whereas prior epidemiological work has established a phenotypic association between stress-related disorders and subsequent autoimmune diseases, including hypothyroidism (Song et al., 2018), our results are the first, to the best of our knowledge, to identify a significant genetic association between hypothyroidism and PTSD. A phenome-wide genetic correlation analysis of thyroid disorders showed that hypothyroidism is genetically correlated with several behavioral traits including fatigue, anxiety, depression, loneliness, and mood swings (Ravera et al., 2018). It remains to be determined whether these associations are causal, and whether any therapeutic implications — for hypothyroidism, PTSD, or other psychiatric disorders — can be drawn from these observations. CMap indicates that *PLEKHM1* has several drug re-purposing candidates (dopamine receptor antagonists, acetylcholine receptor antagonists, and angiotensin receptor antagonists) that could be further investigated in this respect.

With further consideration to drug repurposing opportunities, the possible utility of a dopamine receptor antagonist is intriguing given our prior findings of an association of PTSD re-experiencing symptoms with midbrain medium spiny neurons (Gelernter et al., 2019) which express D1, D2, or both receptors, in conjunction with clinical findings that dopamine (D2) receptor antagonists may benefit some patients with PTSD (Villarreal et al., 2016). On the other hand, a growing body of preclinical evidence points to a possible role for dopamine *augmentation* in fear extinction (Kalisch et al., 2019; Luo et al., 2018; McCullough et al., 2018), which might imply that drugs that *increase* dopamine signaling may have therapeutic potential; these data are supported by our preliminary results showing benefits of methylphenidate for PTSD (McAllister et al., 2016). Interindividual variation may also be relevant in determining treatment response. There is evidence that repeated stress exposure disrupts D1 signaling in the prefrontal cortex (Shinohara et al., 2018). Given the complexities of dopamine signaling (different receptor subtypes, brain regional and cell-type-specific differences in dopaminergic regulation, temporal-specific effects during learning) (Bamford et al., 2018), additional study of these agents is needed to determine their place in PTSD therapeutics.

Our findings also suggest consideration of several other drug classes as therapeutic candidates for PTSD. For example, acetylcholine receptor antagonists could be considered given the convergence of (1) the PheWAS association via *TRAIP* with Chronic Obstructive Pulmonary Disease (COPD; which can be treated with muscarinic receptor antagonists (Ismaila et al., 2015)), and (2) their association in cMAP with *PLEKHM1*. Angiotensin receptor antagonists, also identified as drug candidates through cMAP, have a strong preclinical rationale for use in PTSD (Marvar et al., 2014; Shekhar, 2014) and are, in fact, currently undergoing testing in a randomized placebo-controlled trial of losartan for PTSD (ClinicalTrials.gov Identifier: NCT02709018P).

Our study has numerous limitations and caveats. Whereas we have elected to prioritize post-GWAS analyses based on the largest and most genetically informative PTSD trait available to us — PCL total score, which is reflective of past-month overall symptom severity — it is not currently known whether genetic risk for PTSD differs by trauma type (e.g., combat exposure vs. civilian trauma exposure) or timing (e.g., childhood maltreatment vs. adult assault). Studies of even larger sample size (which MVP will attain in the coming years) and greater granularity with regard to types and chronology of trauma exposure, will be needed to address these questions. Another limitation is our reliance on the European ancestry sample, which reflects both a relative paucity of African and other ancestry individuals (despite this study having the largest African ancestry sample in any PTSD study to date) as well as limitations in the tools available (e.g., LDSC) to conduct post-GWAS analyses in non-European ancestry samples. We must also acknowledge that the drug repurposing propositions, while hypothesis-generating and intriguing, are just that. They are one piece of information that would increase interest in testing the proposed drug classes in patients with PTSD; they must be buttressed by additional preclinical models and complementary informatic approaches (Le-Niculescu et al., 2019) supporting their use, as well as serious consideration of their safety in this population. Lastly, we remind the reader that the present analyses rested solely on GWAS, and with a rather sparse genotyping array, thereby limiting inquiry to common genetic variants; whole genome sequencing is currently underway on a subset of this cohort with the role of identifying rare variants. Epigenetic factors may play a role in a disorder such as PTSD, which has traumatic stress as its precursor (Daskalakis et al., 2018). Many other functional genomics tools can and should be brought to bear on the study of PTSD, expanding the scope of inquiry to encompass a holistic, integrative functional genomic analysis (Li et al., 2018) of this common, serious, and yet still poorly understood neuropsychiatric disorder.

## SUMMARY OF METHODS

### -- EXPERIMENTAL MODEL AND SUBJECT DETAILS

Subjects: All subjects are enrollees in the VA Million Veteran Program (MVP) (Gaziano et al., 2016). Active users of the VHA healthcare system learn of MVP via an invitational mailing and/or through MVP staff while receiving clinical care with informed consent and HIPAA authorization as the only inclusion criteria. As of July 2019, more than 750,000 veterans have enrolled in the program; for the current analyses, genotype data were available from approximately 375,000 participants. Individuals with EHR diagnoses of schizophrenia or bipolar disorder were excluded from participation in this study of PTSD. Research involving MVP is approved by the VA Central IRB; the current project was also approved by VA IRBs in Boston, San Diego, and West Haven.

#### PTSD Case-Control (Binary) Electronic Health Record Derived Phenotype

Details on the derivation and psychometric properties of this phenotype are included in our recent publication (Harrington et al., 2019). In brief, we used manual chart review (*n* = 500) as the gold standard. For both the algorithm and chart review, three classifications were possible: likely PTSD, possible PTSD, no PTSD. We used Lasso regression with cross-validation first to select statistically significant predictors of PTSD from the electronic health record (EHR) and then to generate a predicted probability score of being a PTSD case for every participant in the study population. Probability scores ranged from 0-1.00. Comparing the performance of our probabilistic approach (Lasso algorithm) to a rule-based approach (ICD algorithm), the Lasso algorithm showed modestly higher overall percent agreement with chart review compared to the ICD algorithm (80% vs. 75%), higher sensitivity (.95 vs. .84), and higher overall accuracy (AUC = .95 vs. 90). For purposes of the case-control binary EHR-derived phenotype used here, we applied a 0.7 probability cut point to the Lasso results to determine final PTSD case and control status; we also selected a threshold score of 30 on the PCL from the MVP survey to minimize false negative classifications (e.g., due to an absence of PTSD screening information in the EHR). This final algorithm had a 0.96 sensitivity, 0.98 specificity, 0.91 positive predictive value, and 0.99 negative predictive value for PTSD classification in the trans-ancestral sample as determined by chart review.

#### PTSD Symptom Severity (Quantitative Trait) Sub-phenotypes

The second optional questionnaire, the MVP Lifestyle Survey, includes the PTSD Symptom Checklist (PCL; DSM-IV version) (Blanchard et al., 1996) which asks respondents to report how much they have been bothered *in the past month* by symptoms in response to “stressful life experiences. The PCL has 17 items, each scored on a 5-point severity scale (1 = “Not at All” though 5 = “Extremely”). The re-experiencing (REX) symptom domain is covered by 5 items (score range 5-25), the avoidance (AVOID) domain by 7 items (score range 5-35), and the hyperarousal (HYPER) domain by 5 items (score range 5-25), yielding an overall severity score (TOTAL) for the 17 items (score range 17-85). PCL items and their distributions in EAs and AAs are shown in Table 1. After accounting for missing phenotype data, the final sample size for TOTAL was 186,689 in the EUR sample and 25,318 in the AFR sample.

#### -- Genotyping, Imputation and Quality Control

Genotyping, imputation, and quality control within MVP has been previously described (Gaziano et al., 2016). Briefly, samples were genotyped using a 723,305-SNP Affymetrix Axiom biobank array, customized for MVP. Imputation was performed with minimac3 (Das et al., 2016) using data from the 1000 Genomes project. For post-imputation QC, SNPs with imputation INFO scores of < 0.3 or minor allele frequencies (MAF) below 0.01 were removed from analysis. For the first tranche of data, 22,183 SNPs were selected through linkage disequilibrium (LD) pruning using PLINK (Chang et al., 2015; Purcell et al., 2007) and then Eigensoft (Price et al., 2006) was used to conduct principal component analysis on 343,286 MVP samples and 2,504 1000 Genomes samples (Genomes Project et al., 2015). The reference population groups in the 1000 Genomes samples were used to define EUR (n=241,541) and AFR (n=61,796) groups used in these analyses. Similar methods were used in the second tranche of data, which contained 108,416 new MVP samples and the same 2,504 1000 Genomes samples. In Tranche 2, 80,694 participants were defined as EUR and 20,584 were defined as AFR. In this manuscript, we report results as the meta-analysis of Tranche 1 and 2 data, either for EUR and AFR separately, or as a trans-ancestral meta-analysis.

### -- QUANTIFICATION AND STATISTICAL ANALYSIS

#### Association Analyses

GWAS analysis was carried out by logistic (for the two binary traits) or linear (for the quantitative traits) regression for each ancestry group and tranche using PLINK 2.0 on dosage data, covarying for age, sex, and the first 10 PCs. Meta-analysis was performed using METAL.

#### LD Score Regression (LDSC) and SNP-based Heritability

We used LD score regression (LDSC) through LD Hub (Zheng et al., 2017) to estimate SNP-based heritability, and to assess genetic correlation of PTSD binary algorithmic diagnosis and total PTSD symptoms severity with other traits in LD Hub. LDSC results for 232 traits were extracted from the data at LDHub v2.0 (http://ldsc.broadinstitute.org/ldhub/).

#### Conditional Analysis for Major Depression and other Psychiatric Disorders

Considering the extensive comorbidity between major depression and PTSD (Koenen et al., 2008) we conducted conditional analysis with mtCOJO (Zhu et al., 2018) using GCTA software with the MVP PCL-total symptom severity summary statistics as the primary analysis and the PGC MDD2 (excluding 23andMe due to data unavailability) (Wray et al., 2018) summary statistics to condition the analysis for depression. Additional summary statistics for autism spectrum disorder, anorexia nervosa, anxiety (case-control), alcohol dependence, schizophrenia, bipolar disorder, and attention deficit hyperactivity disorder were obtained from https://www.med.unc.edu/pgc/results-and-downloads/.

#### Gene-based Tests

Summary statistics from the GWAS were loaded into Functional Mapping and Annotation of Genome-Wide Association Studies (FUMA GWAS) (de Leeuw et al., 2015) to test for gene-level associations using Multi-Marker Analysis of GenoMic Annotation (MAGMA) (Watanabe et al., 2017). Input SNPs were mapped to 17,927 protein coding genes. The GWS threshold for the gene-based test was therefore determined to be p = 0.05/17,927 = 2.79×10^−6^.

#### Polygenic Risk Score (PRS) Analysis

Summary statistics from the MVP PTSD EHR—derived binary algorithmic and PCL-total symptom severity analyses, respectively, were used as the base data for calculating PRSs (using PRSice v 1.25) (Euesden et al., 2015).

#### Postmortem Brain Analyses

Quantitative real-time PCR (qRT-PCR) was performed on dissected tissue from Brodmann Area 25, the subgenual prefrontal cortex (sgPFC) in 22 subjects with PTSD (mean age, 46 +/− 11.7 years, 11 females) and 22 matched controls (mean age, 47.8 +/− 11.7 years, 11 females). At the time of death 15 of 22 subjects with PTSD were on an antidepressant and 13 of 22 control subjects were on an over the counter medication. The average postmortem interval (PMI) was 16.1 hours in the PTSD cohort and 18.8 hours for the controls. There were no significant differences between the PTSD and control samples in age, PMI, pH, or RNA integrity number.

qRT-PCR was performed using primers designed to detect the transcript of *CRHR1*. mRNA was isolated from the sgPFC using the RNEasy Plus Mini Kit (Qiagen, Venlo, Netherlands); 1 ug of mRNA was reverse-transcribed into cDNA using the iScript cDNA Synthesis kit (Bio-Rad, Hercules, CA). RNA was hydrolyzed and resuspended in nuclease free water. Gene specific primers for *CRHR1*(Fwd: TGGATGTTCATCTGCATTGG; Rev: GGCCCTGGTAGATGTAGTCG) and the control gene *GAPDH* (Fwd: ACCCAGAAGACTGTGGATGG; Rev: GAGGCAGGGATGATGTTCTG) were designed using Primer 3 v.0.4.0 freeware (http://bioinfo.ut.ee/primer3-0.4.0/) and tested for efficiency and specificity by serial dilution and melt curve analysis. Sybr Green mix (Bio-Rad, Hercules, CA) was used to amplify cDNA. Fold regulation was calculated by using the -delta delta Ct (2-DDCt) method. The 2-DDCt analysis calculates relative gene expression levels between two samples by using the threshold cycles calculated by increasing fluorescent signal of the amplicon during PCR.

For the postmortem gene expression analysis, differences in transcriptional changes between PTSD and controls were evaluated using Graphpad Prism v7 (Graphpad Software, San Diego, CA) with fold changes calculated using the 2-DDCt method. Mann Whitney U test followed by Bonferroni correction for multiple comparisons were used to assess statistical differences in the fold regulation calculated.

#### PrediXcan-S Methods

To perform transcriptome wide association analysis, PrediXcan-S (also known as MetaXcan) (Barbeira et al., 2018) was used to impute gene expression based on the GWAS summary statistics (meta-analysis of tranche 1 and 2 EAs) of PCL-17 Total with the reference gene expression data of 50 tissues from GTEx V7. Gene-expression association with PTSD PCL-17 Total was performed for each tissue (13 of which are brain tissues) individually.

#### Phenome-Wide Association Study (PheWAS)

To identify pleiotropic effects of variants that were significantly associated with PCL-sum score in Europeans, we tested associations between the top 15 loci (Table 4) and every other available phenotype, i.e., a phenome-wide association study (PheWAS) in the MVP sample for each variant.

Binary phenotypes were constructed from manually curated groups of ICD-9 codes (or phecodes), as defined by Denny and colleagues (Denny et al., 2013; Denny et al., 2010). Following the approach of another PheWAS conducted within the MVP (Cai et al., 2018), veterans were defined as a case if they had ≥2 phecodes, very rare phenotypes (prevalence ≤ 0.1%) were excluded, and any relatives greater than 3^rd^ degree (φ ≥ 1/16) were excluded (Manichaikul et al., 2010). A logistic regression was used to measure and test the association between each individual variant with each phenotype. All models were adjusted for age, sex, 20 principal components (PC) to account for population stratification (Cook and Morris, 2016), follow-up time in (log of) months, and (log of) total number of ICD-9 codes as a proxy for healthcare utilization. Association p-values were Bonferroni-corrected to control familywise error rate (FWER) at 5% within each PheWAS; associations with p-values ≤ 0.05/(1335 phenotypes) = 3.75×10^−5^ were considered significant.

#### Drug Repositioning Analysis

CMap (https://clue.io/cmap) provides expression similarity scores for a specific expression profile with other drug-induced transcriptional profiles, including consensus transcriptional signatures of 83 drug classes, i.e., transcriptional profiles induced by 2,837 drugs grouped into 83 drug classes. Expression similarity is evaluated by means of scores that vary from −100 to 100, with -100 the most extreme opposite expression profile and 100 the most extreme similar expression profile.

## CONSORTIA

Names of all consortium members are available from the authors upon request.

## ACKNOWLEDGMENTS

This research is based on data from the Million Veteran Program (MVP), Department of Veterans Affairs Office of Research and Development, and was supported by MVP and the VA Cooperative Studies Program (CSP) study #575B.

## AUTHOR CONTRIBUTIONS

M.B.S., J.G., J.C., K.R., and M.A. had primary responsibility for design and supervision of the study, and managed and organized the group. D.F.L., Z.C., F.R.W., and R.A. contributed to genetic and bioinformatic analyses. K.H., K.C., R.Q., Y-N.A.H., K.R., M.A. and D.P. contributed to phenotyping and phenomic analyses. M.J.G. and R.S.D. conducted the postmortem brain analyses. Initial manuscript was drafted by M.B.S., D.F.L., R.P. and J.G. Manuscript contributions and interpretation of results were provided by M.B.S., D.F.L. Z.C., F.R.W., K.H., M.J.G., D.P., R.S.D, H.Z., R.P., J.C. and J.G. The remaining authors contributed to other organizational or data processing components of the study. All authors saw, had the opportunity to comment on, and approved the final draft.

## DISCLAIMER

The views expressed in this article are those of the authors and do not necessarily reflect the position or policy of the Department of Veterans Affairs or the United States government.

