## Supplemental Table 1 for "Genomic Characterization of Posttraumatic Stress Disorder in a Large US Military Veteran Sample"

Supplemental Table 1: 77 Statistically Significant Genes for European PCL-Total GWGAS

| Gene ID | CHR | START | STOP | NSNPS | NPARAM | N | ZSTAT | P | Gene Symbol |
|---|---|---|---|---|---|---|---|---|---|
| 7473 | 17 | 44839872 | 44896126 | 123 | 23 | 140000 | 7.0247 | 1.07E-12 | WNT3 |
| 246744 | 17 | 44076616 | 44077060 | 1 | 1 | 140000 | 6.7935 | 5.47E-12 | STH |
| 9842 | 17 | 43513266 | 43568146 | 148 | 7 | 140000 | 6.6576 | 1.39E-11 | PLEKHM1 |
| 201176 | 17 | 43471268 | 43510282 | 102 | 7 | 140000 | 6.6176 | 1.82E-11 | ARHGAP27 |
| 100506084 | 17 | 44351550 | 44439416 | 212 | 8 | 140000 | 6.5465 | 2.95E-11 | ARL17B |
| 4905 | 17 | 44668035 | 44834830 | 98 | 11 | 140000 | 6.4607 | 5.21E-11 | NSF |
| 1394 | 17 | 43697710 | 43913194 | 1056 | 10 | 140000 | 6.4284 | 6.45E-11 | CRHR1 |
| 9884 | 17 | 44316744 | 44415160 | 489 | 7 | 140000 | 6.405 | 7.52E-11 | LRRC37A |
| 4137 | 17 | 43971702 | 44105700 | 760 | 5 | 140000 | 6.2992 | 1.50E-10 | MAPT |
| 284058 | 17 | 44107282 | 44302740 | 1055 | 5 | 140000 | 6.2621 | 1.90E-10 | KANSL1 |
| 162540 | 17 | 43922256 | 43924438 | 17 | 2 | 140000 | 6.2313 | 2.31E-10 | SPPL2C |
| 4987 | 20 | 62711451 | 62731996 | 69 | 22 | 140000 | 5.9584 | 1.27E-09 | OPRL1 |
| 9807 | 3 | 49761728 | 49823973 | 108 | 12 | 140000 | 5.9187 | 1.62E-09 | IP6K1 |
| 63891 | 3 | 49726950 | 49758962 | 51 | 10 | 140000 | 5.8559 | 2.37E-09 | RNF123 |
| 79012 | 3 | 49895414 | 49907655 | 18 | 5 | 140000 | 5.6667 | 7.28E-09 | CAMKV |
| 51326 | 17 | 44577184 | 44657155 | 37 | 7 | 140000 | 5.6598 | 7.58E-09 | ARL17A |
| 474170 | 17 | 44587323 | 44633016 | 27 | 6 | 140000 | 5.6337 | 8.82E-09 | LRRC37A2 |
| 10293 | 3 | 49866028 | 49893992 | 42 | 7 | 140000 | 5.5451 | 1.47E-08 | TRAIP |
| 64839 | 5 | 1.07E+08 | 1.08E+08 | 1238 | 46 | 140000 | 5.5398 | 1.51E-08 | FBXL17 |
| 93986 | 7 | 1.14E+08 | 1.14E+08 | 1038 | 53 | 140000 | 5.5239 | 1.66E-08 | FOXP2 |
| 389118 | 3 | 49828165 | 49837254 | 17 | 7 | 140000 | 5.5033 | 1.86E-08 | CDHR4 |
| 84315 | 3 | 49946302 | 49967445 | 28 | 9 | 140000 | 5.4726 | 2.22E-08 | MON1A |
| 56145 | 5 | 1.4E+08 | 1.4E+08 | 455 | 25 | 140000 | 5.4651 | 2.31E-08 | PCDHA3 |
| 4486 | 3 | 49924435 | 49941311 | 23 | 4 | 140000 | 5.4125 | 3.11E-08 | MST1R |
| 8379 | 7 | 1855428 | 2272583 | 1933 | 38 | 140000 | 5.3429 | 4.57E-08 | MAD1L1 |
| 10180 | 3 | 49977474 | 50114685 | 266 | 13 | 140000 | 5.3281 | 4.96E-08 | RBM6 |
| 10669 | 2 | 27322159 | 27341995 | 31 | 6 | 140000 | 5.3152 | 5.33E-08 | CGREF1 |
| 8555 | 9 | 99252807 | 99382112 | 113 | 19 | 140000 | 5.3147 | 5.34E-08 | CDC14B |
| 56144 | 5 | 1.4E+08 | 1.4E+08 | 432 | 23 | 140000 | 5.2791 | 6.49E-08 | PCDHA4 |
| 10181 | 3 | 50126341 | 50156397 | 29 | 6 | 140000 | 5.2501 | 7.60E-08 | RBM5 |
| 203062 | 8 | 1.43E+08 | 1.43E+08 | 795 | 39 | 140000 | 5.1817 | 1.10E-07 | TSNARE1 |
| 56141 | 5 | 1.4E+08 | 1.4E+08 | 382 | 21 | 140000 | 5.164 | 1.21E-07 | PCDHA7 |
| 6925 | 18 | 52889562 | 53303252 | 691 | 57 | 140000 | 5.1028 | 1.67E-07 | TCF4 |
| 56143 | 5 | 1.4E+08 | 1.4E+08 | 398 | 20 | 140000 | 5.0786 | 1.90E-07 | PCDHA5 |
| 56142 | 5 | 1.4E+08 | 1.4E+08 | 393 | 20 | 140000 | 5.0639 | 2.05E-07 | PCDHA6 |
| 27086 | 3 | 71003865 | 71633140 | 1365 | 123 | 140000 | 5.0597 | 2.10E-07 | FOXP1 |
| 22927 | 9 | 99212414 | 99253618 | 83 | 9 | 140000 | 5.0172 | 2.62E-07 | HABP4 |
| 169792 | 9 | 3824128 | 4300036 | 2094 | 165 | 140000 | 5.0075 | 2.76E-07 | GLIS3 |
| 56146 | 5 | 1.4E+08 | 1.4E+08 | 466 | 22 | 140000 | 4.9727 | 3.30E-07 | PCDHA2 |
| 9752 | 5 | 1.4E+08 | 1.4E+08 | 362 | 20 | 140000 | 4.9726 | 3.30E-07 | PCDHA9 |
| 56896 | 2 | 27070969 | 27173219 | 156 | 11 | 140000 | 4.9707 | 3.34E-07 | DPYSL5 |
| 4690 | 3 | 1.37E+08 | 1.37E+08 | 199 | 7 | 140000 | 4.947 | 3.77E-07 | NCK1 |
| 5789 | 9 | 8314246 | 10612723 | 10785 | 491 | 140000 | 4.9301 | 4.11E-07 | PTPRD |
| 56147 | 5 | 1.4E+08 | 1.4E+08 | 483 | 21 | 140000 | 4.9191 | 4.35E-07 | PCDHA1 |
| 56140 | 5 | 1.4E+08 | 1.4E+08 | 373 | 19 | 140000 | 4.8995 | 4.80E-07 | PCDHA8 |
| 22864 | 12 | 57647547 | 57824782 | 223 | 15 | 140000 | 4.8934 | 4.95E-07 | R3HDM2 |
| 10677 | 12 | 58190415 | 58210193 | 24 | 9 | 140000 | 4.8832 | 5.22E-07 | AVIL |
| 25789 | 19 | 18723682 | 18731849 | 34 | 10 | 140000 | 4.8612 | 5.83E-07 | TMEM59L |
| 2186 | 17 | 65821644 | 65980494 | 297 | 14 | 140000 | 4.8386 | 6.54E-07 | BPTF |
| 23224 | 14 | 64319683 | 64693167 | 1013 | 50 | 140000 | 4.8132 | 7.43E-07 | SYNE2 |
| 5188 | 4 | 1.53E+08 | 1.53E+08 | 226 | 12 | 140000 | 4.7974 | 8.04E-07 | GATB |
| 3626 | 12 | 57828543 | 57844609 | 27 | 5 | 140000 | 4.7887 | 8.39E-07 | INHBC |
| 406 | 11 | 13299325 | 13408813 | 265 | 36 | 140000 | 4.7581 | 9.77E-07 | ARNTL |
| 55049 | 19 | 18699495 | 18703147 | 11 | 3 | 140000 | 4.7567 | 9.84E-07 | C19orf60 |
| 22924 | 2 | 27193525 | 27250087 | 59 | 5 | 140000 | 4.7464 | 1.04E-06 | MAPRE3 |
| 6671 | 7 | 21467689 | 21554440 | 266 | 25 | 140000 | 4.7315 | 1.11E-06 | SP4 |
| 56136 | 5 | 1.4E+08 | 1.4E+08 | 292 | 17 | 140000 | 4.7212 | 1.17E-06 | PCDHA13 |
| 60509 | 2 | 27274291 | 27293490 | 24 | 5 | 140000 | 4.7199 | 1.18E-06 | AGBL5 |
| 6405 | 3 | 50192562 | 50226508 | 65 | 11 | 140000 | 4.7113 | 1.23E-06 | SEMA3F |
| 56139 | 5 | 1.4E+08 | 1.4E+08 | 340 | 18 | 140000 | 4.707 | 1.26E-06 | PCDHA10 |
| 56135 | 5 | 1.4E+08 | 1.4E+08 | 212 | 18 | 140000 | 4.7069 | 1.26E-06 | PCDHAC1 |
| 1496 | 2 | 79740060 | 80875993 | 3464 | 236 | 140000 | 4.7021 | 1.29E-06 | CTNNA2 |
| 23438 | 5 | 1.4E+08 | 1.4E+08 | 11 | 3 | 140000 | 4.6949 | 1.33E-06 | HARS2 |
| 55843 | 2 | 1.44E+08 | 1.45E+08 | 1594 | 91 | 140000 | 4.6897 | 1.37E-06 | ARHGAP15 |
| 51075 | 11 | 57479995 | 57508445 | 46 | 6 | 140000 | 4.6605 | 1.58E-06 | TMX2 |
| 23163 | 17 | 73232694 | 73258444 | 50 | 8 | 140000 | 4.6484 | 1.67E-06 | GGA3 |
| 3795 | 2 | 27309611 | 27323619 | 14 | 2 | 140000 | 4.6324 | 1.81E-06 | KHK |
| 54932 | 9 | 1.4E+08 | 1.4E+08 | 379 | 64 | 140000 | 4.6295 | 1.83E-06 | EXD3 |
| 9244 | 19 | 18704035 | 18717660 | 46 | 8 | 140000 | 4.6077 | 2.04E-06 | CRLF1 |
| 56137 | 5 | 1.4E+08 | 1.4E+08 | 301 | 17 | 140000 | 4.5902 | 2.21E-06 | PCDHA12 |
| 10287 | 20 | 62704534 | 62711356 | 13 | 5 | 140000 | 4.5898 | 2.22E-06 | RGS19 |
| 7187 | 14 | 1.03E+08 | 1.03E+08 | 283 | 17 | 140000 | 4.5812 | 2.31E-06 | TRAF3 |
| 56138 | 5 | 1.4E+08 | 1.4E+08 | 312 | 16 | 140000 | 4.5643 | 2.51E-06 | PCDHA11 |
| 3035 | 5 | 1.4E+08 | 1.4E+08 | 30 | 5 | 140000 | 4.5546 | 2.62E-06 | HARS |
| 55374 | 5 | 1.4E+08 | 1.4E+08 | 11 | 3 | 140000 | 4.5525 | 2.65E-06 | TMCO6 |
| 9702 | 11 | 95523625 | 95565857 | 148 | 14 | 140000 | 4.5418 | 2.79E-06 | CEP57 |
| 5253 | 9 | 96338909 | 96441869 | 469 | 27 | 140000 | 4.5173 | 3.13E-06 | PHF2 |
