## Supplemental Tables 2a and 2b for "Genomic Characterization of Posttraumatic Stress Disorder in a Large US Military Veteran Sample"

**Supplemental Table 2a.** Linkage disequilibrium score regression results from the MVP EUR Total PCL summary statistics. Red and blue indicates positive or negative genetic correlation with Total PCL, respectively. P-values < 0.0001 are considered statistically significant, taking into account the number of tests conducted (Bonferroni correction)

| Trait | PubMed ID | rg | se | z | P-value |
|---|---|---|---|---|---|
| Depressive symptoms | 27089181 | 0.7412 | 0.0427 | 17.3422 | 2.26E-67 |
| Insomnia | 28604731 | 0.4886 | 0.0537 | 9.0986 | 9.15E-20 |
| Schizophrenia | 25056061 | 0.234 | 0.0285 | 8.2209 | 2.02E-16 |
| Smoking Initiation | 20418890 | 0.4064 | 0.0499 | 8.1516 | 3.59E-16 |
| Neuroticism | 24828478 | 0.6692 | 0.0957 | 6.995 | 2.65E-12 |
| Waist-to-hip ratio | 25673412 | 0.2117 | 0.032 | 6.6084 | 3.88E-11 |
| Number of children ever born | 27798627 | 0.2945 | 0.0461 | 6.3947 | 1.61E-10 |
| Waist circumference | 25673412 | 0.1704 | 0.0311 | 5.478 | 4.30E-08 |
| Coronary artery disease | 26343387 | 0.1715 | 0.0317 | 5.4083 | 6.36E-08 |
| Obesity class 3 | 23563607 | 0.328 | 0.061 | 5.3787 | 7.50E-08 |
| Obesity class 1 | 23563607 | 0.182 | 0.0349 | 5.21 | 1.89E-07 |
| Body fat | 26833246 | 0.2317 | 0.0463 | 5 | 5.73E-07 |
| Body mass index | 20935630 | 0.154 | 0.0316 | 4.8658 | 1.14E-06 |
| PGC cross-disorder analysis | 23453885 | 0.2167 | 0.0456 | 4.7521 | 2.01E-06 |
| Obesity class 2 | 23563607 | 0.1867 | 0.0417 | 4.4789 | 7.50E-06 |
| Lung cancer | 24880342 | 0.3593 | 0.0832 | 4.3201 | 1.56E-05 |
| ADHD | 27663945 | 0.5106 | 0.1187 | 4.3018 | 1.69E-05 |
| Triglycerides | 20686565 | 0.1515 | 0.0376 | 4.0341 | 5.48E-05 |
| Rheumatoid Arthritis | 24390342 | 0.1706 | 0.0433 | 3.9414 | 8.10E-05 |
| Extreme bmi | 23563607 | 0.1862 | 0.048 | 3.8749 | 0.0001 |
| Parents age at death | 27015805 | -0.3347 | 0.0805 | -4.158 | 3.21E-05 |
| Mothers age at death | 27015805 | -0.3171 | 0.0605 | -5.2384 | 1.62E-07 |
| Smoking Cessation | 20418890 | -0.4342 | 0.0781 | -5.5594 | 2.71E-08 |
| Childhood IQ | 23358156 | -0.4679 | 0.0836 | -5.5961 | 2.19E-08 |
| Fathers age at death | 27015805 | -0.4396 | 0.0757 | -5.8078 | 6.33E-09 |
| Subjective well being | 27089181 | -0.3934 | 0.0418 | -9.4114 | 4.90E-21 |
| College completion | 23722424 | -0.5096 | 0.0459 | -11.0997 | 1.26E-28 |
| Intelligence | 28530673 | -0.4595 | 0.0378 | -12.157 | 5.26E-34 |
| Age of first birth | 27798627 | -0.5273 | 0.0372 | -14.1604 | 1.61E-45 |
| Years of schooling 2016 | 27225129 | -0.4223 | 0.0264 | -16.0063 | 1.16E-57 |

**Supplemental Table 2b.** Linkage disequilibrium score regression results from the MVP Algorithmic Case-Control summary statistics. Red and blue indicates positive or negative genetic correlation with Case-Control status, respectively. P-values < 0.0001 are considered statistically significant, taking into account the number of tests conducted (Bonferroni correction)

| Trait | PubMed ID | rg | se | z | P-value |
|---|---|---|---|---|---|
| Depressive symptoms | 27089181 | 0.553 | 0.0598 | 9.2391 | 2.48E-20 |
| Neuroticism | 27089181 | 0.4173 | 0.047 | 8.8777 | 6.83E-19 |
| Smoking Initiation | 20418890 | 0.403 | 0.0623 | 6.4635 | 1.02E-10 |
| Number of children ever born | 27798627 | 0.3023 | 0.0509 | 5.9411 | 2.83E-09 |
| Insomnia | 28604731 | 0.38 | 0.0643 | 5.908 | 3.46E-09 |
| Coronary artery disease | 26343387 | 0.1804 | 0.041 | 4.3973 | 1.10E-05 |
| Schizophrenia | 25056061 | 0.1749 | 0.0411 | 4.2577 | 2.07E-05 |
| Obesity class 3 | 23563607 | 0.3166 | 0.0744 | 4.2566 | 2.08E-05 |
| Waist-to-hip ratio | 25673412 | 0.1619 | 0.0386 | 4.1977 | 2.70E-05 |
| Waist circumference | 25673412 | 0.149 | 0.0377 | 3.9507 | 7.79E-05 |
| Parents age at death | 27015805 | -0.3819 | 0.0999 | -3.8239 | 0.0001 |
| Smoking Cessation | 20418890 | -0.4427 | 0.0952 | -4.6493 | 3.33E-06 |
| Childhood IQ | 23358156 | -0.3925 | 0.0839 | -4.6788 | 2.89E-06 |
| Fathers age at death | 27015805 | -0.3904 | 0.079 | -4.9443 | 7.64E-07 |
| Intelligence | 28530673 | -0.4274 | 0.0469 | -9.1158 | 7.81E-20 |
| College completion | 23722424 | -0.5493 | 0.0528 | -10.4122 | 2.18E-25 |
| Age of first birth | 27798627 | -0.5307 | 0.0422 | -12.5631 | 3.37E-36 |
| Years of schooling | 27225129 | -0.4217 | 0.0314 | -13.4168 | 4.82E-41 |
