## Supplemental Table 3 for "Genomic Characterization of Posttraumatic Stress Disorder in a Large US Military Veteran Sample"

### Supplemental Table 3. Replication Results of MVP SNPs into PGC-PTSD 2.0

| | | | | MVP | | | PGC | | | | | LEGEND |
|---|---|---|---|---|---|---|---|---|---|---|---|---|
| | MarkerName | Allele1 | Allele2 | BETA | SE | P | Effect | StdErr | P-value | Ncases | Ncontrols | nominal replication |
| EUR-case-controlPTSD | rs10767744 | t | c | -0.0562 | 0.0088 | 1.75E-10 | -0.0006 | 0.0153 | 0.9693 | 23185 | 151309 | direction replication |
| | rs137999048 | cca | c | 0.1159 | 0.0202 | 1.03E-08 | -0.0177 | 0.0403 | 0.6613 | 12498 | 33851 | |
| | rs7680 | a | g | -0.0712 | 0.013 | 4.17E-08 | -0.0639 | 0.0215 | 0.003012 | 23185 | 151309 | |
| EUR-PCLtotal | rs55925547 | C | T | 0.4402 | 0.0600 | 2.16E-13 | 0.0505 | 0.0189 | 0.00753 | 22738 | 151098 | |
| | rs10235664 | C | T | -0.3667 | 0.0546 | 1.82E-11 | -0.037 | 0.0169 | 0.02865 | 23185 | 151309 | |
| | rs111488606 | CA | C | 0.3102 | 0.0515 | 1.72E-09 | -0.0019 | 0.0177 | 0.9152 | 12051 | 33640 | |
| | rs13262595 | G | A | -0.2823 | 0.0472 | 2.20E-09 | -0.0326 | 0.0147 | 0.02688 | 23185 | 151309 | |
| | rs2314662 | C | T | -0.3614 | 0.0613 | 3.78E-09 | 0.0096 | 0.0185 | 0.6049 | 23185 | 151309 | |
| | rs10171148 | A | C | 0.2811 | 0.0483 | 5.87E-09 | 0.0527 | 0.0149 | 0.0004196 | 23185 | 151309 | |
| | rs62465629 | C | T | -0.3929 | 0.0676 | 6.30E-09 | -0.0234 | 0.0198 | 0.2387 | 23185 | 151309 | |
| | rs1496246 | G | A | 0.2973 | 0.0513 | 6.60E-09 | -0.011 | 0.0156 | 0.4786 | 23185 | 151309 | |
| | rs251350 | C | T | -0.2538 | 0.0443 | 1.03E-08 | -0.0096 | 0.0165 | 0.5604 | 12823 | 35648 | |
| | rs11507683 | T | C | 0.4137 | 0.0725 | 1.15E-08 | 0.0493 | 0.0238 | 0.03834 | 21980 | 147088 | |
| | rs599550 | A | G | 0.3948 | 0.0692 | 1.18E-08 | 0.029 | 0.0213 | 0.1736 | 23185 | 151309 | |
| | rs4364183 | A | G | 0.3043 | 0.0534 | 1.22E-08 | 0.0146 | 0.0163 | 0.3715 | 23185 | 151309 | |
| | rs10255943 | G | A | 0.3011 | 0.0537 | 2.00E-08 | 0.0442 | 0.0166 | 0.007774 | 23185 | 151309 | |
| | rs62417832 | T | G | 0.2922 | 0.0527 | 2.90E-08 | 0.0123 | 0.0177 | 0.4876 | 23185 | 151309 | |
| | rs111950471 | TATTA | T | -0.2769 | 0.0506 | 4.34E-08 | -0.0237 | 0.0174 | 0.1727 | 12498 | 33851 | |
| EUR-HYPERAROUSAL | rs377112142 (proxy:rs62060852) | CT | C | 0.1323 | 0.0181 | 3.06E-13 | 0.0441 | 0.018 | 0.01418 | 22413 | 149301 | |
| | rs55789728 | A | G | 0.1303 | 0.018 | 4.62E-13 | 0.038 | 0.0177 | 0.03133 | 23185 | 151309 | |
| | rs10992804 (proxy:rs576430065) | CA | C | -0.1206 | 0.0179 | 1.67E-11 | -0.04 | 0.0152 | 0.008671 | 23185 | 151309 | |
| | rs1496246 | A | G | -0.1037 | 0.0163 | 1.77E-10 | 0.011 | 0.0156 | 0.4786 | 23185 | 151309 | |
| | rs547649546 (proxy:rs2271961) | CA | C | -0.0937 | 0.0155 | 1.59E-09 | -0.0107 | 0.0149 | 0.4719 | 22738 | 151098 | |
| | rs2887882 | T | C | -0.1118 | 0.0186 | 1.89E-09 | -0.0255 | 0.0185 | 0.1684 | 23185 | 151309 | |
| | rs7519147 | T | C | -0.0906 | 0.0151 | 1.90E-09 | -0.0176 | 0.0148 | 0.2323 | 23185 | 151309 | |
| | rs13032994 | T | C | 0.0968 | 0.0164 | 3.73E-09 | -0.0193 | 0.0164 | 0.2382 | 23185 | 151309 | |
| | rs113341106 | G | GC | -0.0923 | 0.0157 | 3.82E-09 | -0.0422 | 0.0165 | 0.01069 | 12498 | 33851 | |
| | rs12420134 | C | G | -0.1229 | 0.0212 | 6.45E-09 | -0.0418 | 0.0194 | 0.03059 | 23185 | 151309 | |
| | rs17209774 | C | G | -0.0907 | 0.0157 | 7.97E-09 | 0.016 | 0.0154 | 0.2987 | 23185 | 151309 | |
| | rs11507683 | T | C | 0.131 | 0.023 | 1.16E-08 | 0.0493 | 0.0238 | 0.03834 | 21980 | 147088 | |
| | rs60958094 | T | TATAA | 0.0961 | 0.0171 | 1.99E-08 | 0.0496 | 0.0177 | 0.005102 | 12498 | 33851 | |
| | rs4129585 | A | C | 0.0835 | 0.0149 | 2.07E-08 | 0.032 | 0.0147 | 0.02953 | 23185 | 151309 | |
| | rs7865488 | A | C | -0.1067 | 0.0193 | 3.10E-08 | 0.0038 | 0.0192 | 0.8448 | 23185 | 151309 | |
| | rs201253409 (proxy:rs549326362) | T | G | -0.0884 | 0.0162 | 4.46E-08 | -0.0266 | 0.0174 | 0.1259 | 12498 | 33851 | |
| | rs1156954 | A | G | 0.0896 | 0.0164 | 4.82E-08 | 0.0036 | 0.0157 | 0.8202 | 23185 | 151309 | |
| EUR-AVOID | rs55925547 | t | c | -0.1932 | 0.0263 | 2.08E-13 | -0.0505 | 0.0189 | 0.00753 | 22738 | 151098 | |
| | rs35761884 | ct | c | 0.1388 | 0.0214 | 9.72E-11 | 0.0246 | 0.0165 | 0.1356 | 12498 | 33851 | |
| | rs251350 | t | c | 0.1192 | 0.0194 | 8.15E-10 | 0.0096 | 0.0165 | 0.5604 | 12823 | 35648 | |
| | rs4129585 | a | c | 0.125 | 0.0206 | 1.25E-09 | 0.032 | 0.0147 | 0.02953 | 23185 | 151309 | |
| | rs2314662 | t | c | 0.1599 | 0.0269 | 2.74E-09 | -0.0096 | 0.0185 | 0.6049 | 23185 | 151309 | |
| | rs62465629 | t | c | 0.175 | 0.0296 | 3.54E-09 | 0.0234 | 0.0198 | 0.2387 | 23185 | 151309 | |
| | rs62417832 | t | g | 0.1335 | 0.0231 | 7.04E-09 | 0.0123 | 0.0177 | 0.4876 | 23185 | 151309 | |
| | rs11507683 | t | c | 0.1834 | 0.0318 | 7.74E-09 | 0.0493 | 0.0238 | 0.03834 | 21980 | 147088 | |
| | rs10171148 | a | c | 0.1211 | 0.0212 | 1.07E-08 | 0.0527 | 0.0149 | 0.0004196 | 23185 | 151309 | |
| | rs10235664 | t | c | 0.1337 | 0.0239 | 2.17E-08 | 0.037 | 0.0169 | 0.02865 | 23185 | 151309 | |
| | rs1496246 | a | g | -0.1234 | 0.0224 | 3.66E-08 | 0.011 | 0.0156 | 0.4786 | 23185 | 151309 | |
| EUR-REEXPERIENCING | rs10235664 | t | c | 0.1055 | 0.0169 | 4.66E-10 | 0.037 | 0.0169 | 0.02865 | 23185 | 151309 | |
| | rs242925 | t | c | -0.0931 | 0.015 | 5.50E-10 | -0.0277 | 0.0151 | 0.06757 | 22413 | 149301 | |
| | rs34177209 | a | t | 0.1205 | 0.0216 | 2.34E-08 | 0.0392 | 0.0221 | 0.07641 | 12463 | 33695 | |
| | rs1501485 | a | g | 0.0839 | 0.0147 | 1.22E-08 | 0.0179 | 0.0148 | 0.2259 | 23185 | 151309 | |
| | rs11773880 | t | g | 0.0977 | 0.0174 | 1.97E-08 | 0.0193 | 0.0171 | 0.2583 | 23185 | 151309 | |
| | rs10767734 | t | c | -0.0895 | 0.0157 | 1.23E-08 | 0.0085 | 0.0155 | 0.5824 | 23185 | 151309 | |
| | rs10977193 | a | g | -0.0934 | 0.017 | 4.17E-08 | -0.0048 | 0.0175 | 0.7825 | 23185 | 151309 | |
| | rs2777888 | a | g | -0.0929 | 0.0146 | 2.26E-10 | 0.0026 | 0.0149 | 0.8594 | 22738 | 151098 | |
| | rs35371867 | a | g | 0.1006 | 0.0156 | 1.24E-10 | 0.0018 | 0.0155 | 0.9066 | 23185 | 151309 | |
| | rs6031014 | a | g | -0.3267 | 0.0598 | 4.63E-08 | 0.0069 | 0.066 | 0.9163 | 10602 | 31671 | |
